# Two modes of evolution shape bacterial strain diversity in the gut for thousands of generations

**DOI:** 10.1101/2022.01.11.475860

**Authors:** N. Frazão, A. Konrad, D. Güleresi, M. Lässig, I. Gordo

## Abstract

How and at what pace bacteria evolve when colonizing healthy hosts remains unclear. Here, by monitoring evolution for more than six thousand generations in the mouse gut, we show that the successful colonization of an invader *Escherichia coli* depends on the diversity of the existing microbiota and the presence of a closely related strain. Following colonization, two modes of evolution were observed: one in which diversifying selection leads to long-term coexistence of ecotypes and a second in which directional selection propels selective sweeps. These modes can be quantitatively distinguished by the statistics of mutation trajectories. In our experiments, diversifying selection was marked by the emergence of metabolic mutations, and directional selection by acquisition of prophages, which bring their own benefits and costs. In both modes, we observed parallel evolution, with mutation accumulation rates comparable to those typically observed *in vitro* on similar time scales. Our results show that gut environments can rapidly generate diversifying selection and ecotype formation.

## Main

The well-known capacity of bacteria to adapt to laboratory environments (Behringer et al., 2018; Good et al., 2017; Tenaillon et al., 2016), as well as the high level of evolutionary parallelism that is observed when bacteria are exposed to the same stresses *ex vivo*, demands an understanding of their evolutionary change in natural ecosystems. Time series metagenomics data have started to reveal that significant amounts of evolution can occur in the microbiomes of every human throughout their lifetimes (Garud et al., 2019; Poyet et al., 2019). These genetic changes lead to individualized microbiomes and may provide insight into the main drivers of colonization success for an invading strain or species, with consequences for pathogenesis (Zhao et al., 2019). Microbiome studies allow bacterial evolution to be investigated within hosts and may provide answers to questions such as: What are the selection pressures operating in the gut? How is bacterial evolution influenced by complex communities (Madi et al., 2020; Scheuerl et al., 2020)? How do the different evolutionary mechanisms – mutation, genetic drift, recombination, and natural selection – shape the diversity of the gut ecosystem (Garud et al., 2019)?

Here we use the microbiota of mice to address these questions. Mice can be raised under strict environmental conditions, including a controlled diet, constant temperature and humidity. Thus, we can more easily quantify patterns of bacterial evolution and the effect of a rich microbiome than in humans. Mice can also be used as models for health problems relevant to humans (Peters et al., 2007) and help to understand how microbial traits may cause host phenotypes (Walter et al., 2020).

### Rate of mutation accumulation in invading *Escherichia coli*

We report an *in vivo* long-term experiment that maps the evolution of an invading *Escherichia coli* strain in the presence of a diverse mouse microbiota over thousands of generations. The invasion becomes possible after a short perturbation by an antibiotic (Frazão et al., 2019) (Fig. 1 and Extended Data Fig. 1, Supplementary Tables 1-3). We colonized nine mice with an invading antibiotic resistant *E. coli* carrying a fluorescent marker to track its long-term evolution while resident in the mouse gut (see Methods). In two mice, colonization with the invading *E. coli* was unsuccessful (Fig. 1a, Extended Data Fig. 2, Supplementary Table 2). We found the long-term persistence of the invading species to be negatively correlated with the Shannon index of the mouse microbiota (Log10(CFUs)) (r=−0.78 ((−0.85, −0.67), 95%CI), df=77, p<0.0001), but not with the presence of any resident *E. coli* of a different phylogenetic group (group B1) (Frazão et al., 2019). Interestingly, we found that the resident *E. coli* persists independently of microbiota diversity. This highlights the need to understand diversity at the strain level to predict key functions of the microbiome, such as its capacity to provide colonization resistance to opportunists (Hromada et al., 2021).

**Fig. 1.**
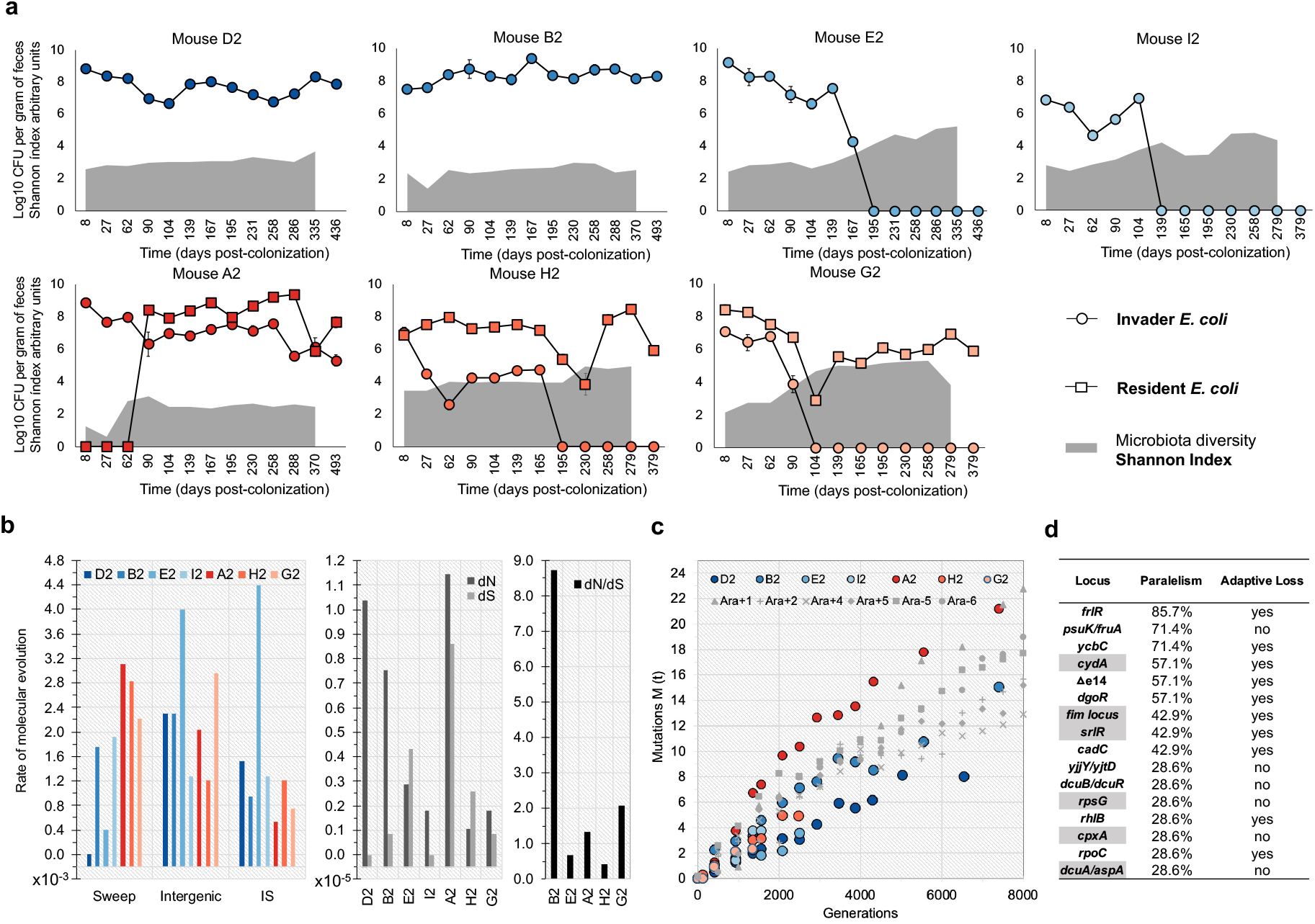
Colonization success depends on microbiota diversity and rates of molecular evolution across mice. **a,** Time series of the abundances of the invader (circles) and resident (squares) *E. coli* lineages and microbiota species diversity (Shannon diversity). The values of microbiota diversity observed are shown as a gray area. Error bars represent 2*Standard Error (2SE). **b,** Rate of molecular evolution, per generation, in each host (sweeps, intergenic mutations, Insertion Sequences). dN and dS, and dN/dS ratio. **c,** *E. coli* mutation accumulation during *in vivo* and *in vitro* evolution (data of non-mutator lines from (Good et al., 2017)). “Mutations M(t)” correspond to the sum of allele frequencies at each sampling point for the *in vitro* and *in vivo* experiments (Supplementary Tables 6 and 8 to 14). Blue and red hues represent mice where the invader colonized the gut alone or together with the *resident E. coli* lineage, respectively. **d,** Level of mutational parallelism in the invader lineage across hosts: the genetic targets of adaptation and the frequency of hosts where it was changed is shown, as well as the changes that likely involve loss of function. Mutation targets also found in Lenski’s *in vitro* evolution experiment, data from^1^, are highlighted in gray.

We followed the long-term evolution of invader *E. coli* (Fig. 1a, Supplementary Table 1) by pool-sequencing of clones sampled from each mouse’s feces. In three mice, long-term colonization (>6000 generations) was achieved, while shorter-term colonization occurred in the other mice. We estimated 15 generations/day for the invader *E. coli* by measuring bacterial ribosomal content (Extended Data Fig. 3 and Supplementary Table 4). Temporal series sequencing allowed estimation of the rate of molecular evolution from the accumulation of novel mutations in the invader *E. coli* genome, isolated from each mouse. Across mice, *E. coli* evolution was characterized by an elevated ratio of non-synonymous to synonymous mutations, indicative of adaptive evolution driven by strong selection (Fig. 1b, Supplementary Table 5). The overall mutation accumulation rate is remarkably similar to Lenski’s long-term evolution of non-mutator *E. coli in vitro* (Good et al., 2017), despite differences in ecology (Fig. 1c and Supplementary Table 6). Across all mice, mutations accumulated at an average rate of 2.1×10^-3^ (0.53×10^-3^ 2SE) per genome per generation, suggestive of a clock-like rate of molecular evolution within a healthy host.

We also detected signs of parallel evolution i.e., changes in the same mutational targets (Tenaillon et al., 2012) within and across mice. The level of parallel evolution varied across gene targets: the locus with the highest parallelism that mutated in *E. coli* populations isolated from six out of seven mice, was *frlR*. In contrast, the locus *dgoR* was mutated only when resident *E. coli* was absent (Fig. 1d, Supplementary Tables 7-14). Evolutionary parallelism was also observed at the insertion sequence (IS) element level, indicating that many of these changes are adaptive (see below and Supplementary Tables 8-14). Across all mice, 277 evolutionary changes were detected (Supplementary Tables 5 and 8 to 14). Of these, 24% (66/277) occurred in intergenic regions, indicating substantial alterations in gene regulation occurred during invader *E. coli* adaptation to the gut. Over the cumulative >29,000 generations of evolution, 46 selective sweeps were observed. We defined a selective sweep as a mutational trajectory that reached >95% frequency and remained at these high frequencies for the duration of observation (Supplementary Table 5). These sweeps were caused by *de novo* substitutions, indels, ISs, and horizontal gene transfer (HGT) events, mediated by phages and plasmids (Supplementary Tables 8-14). Together, our observations show that all invading *E. coli* strains followed fast, adaptive evolutionary dynamics in the gut environment for thousands of generations.

### Diversifying and directional selection

The large-scale dynamics of molecular evolution and its effects on the diversity of the invading strain populations, however, play out in strikingly different ways across the individual mice. Specifically, we observed two modes of evolution: one in which new ecotypes form in the invader *E. coli* and are maintained for >6000 generations (Fig. 2), and a second, in which recurrent selective sweeps intertwined with HGT events (Fig. 3).

**Fig. 2.**
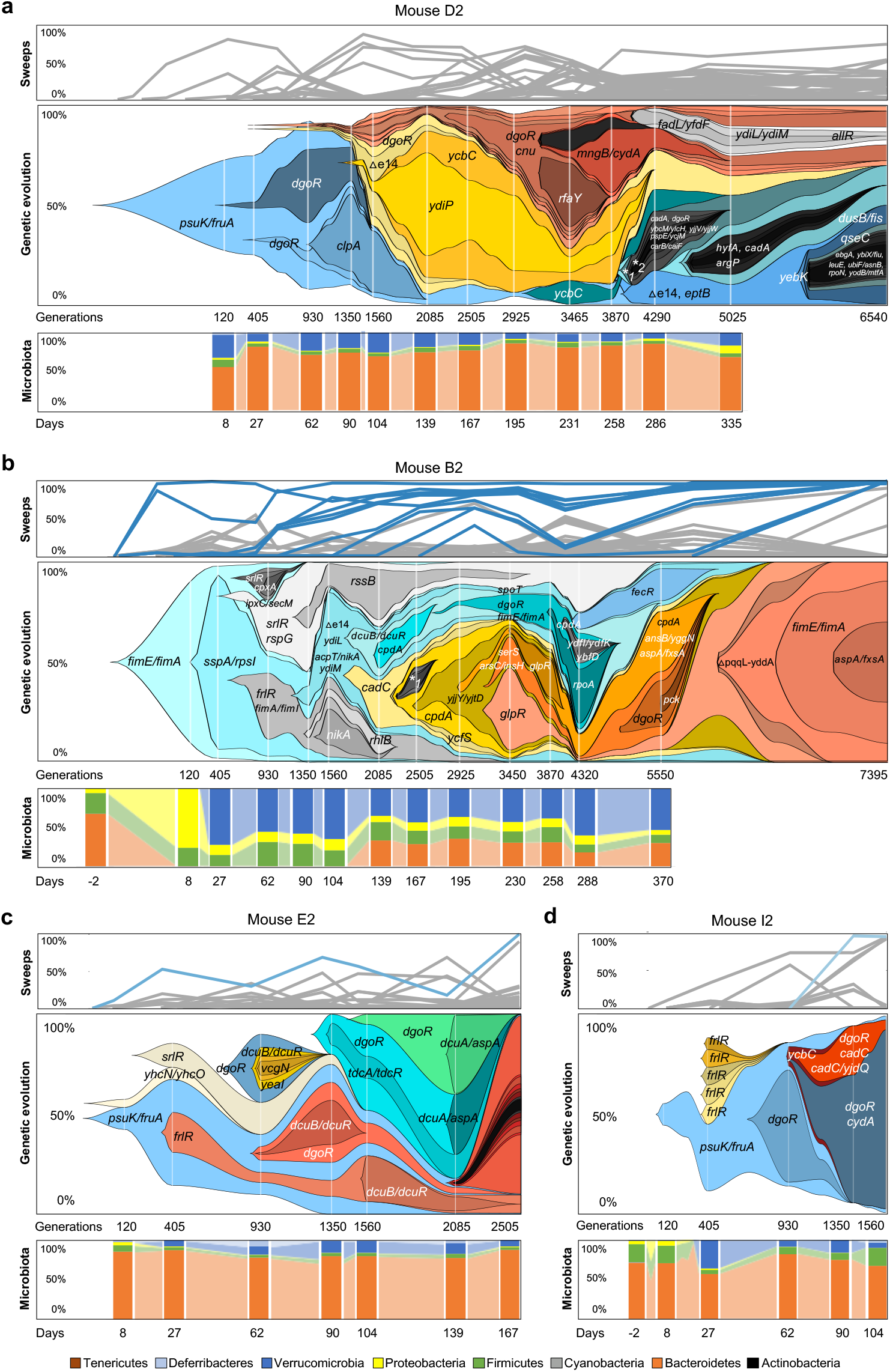
Evolution of invader *E. coli* in the absence of the resident. **a-d,** Frequency of the mutations identified in the invader *E. coli* population across time in mouse D2, B2, E2 and I2 (top panels) and corresponding Muller Plots showing the spread of mutations observed in each mice (middle panels). Microbiota composition (phylum level) along time (bottom panels), showing a high temporal stability in the long term. *1 *fimA, frlR, ykfI, dgoR, dcuA/aspA, qseC, cydA;* *2 *ycbC, rpoC, frlR*. Mutations that reached frequency above 95% are highlighted in color, other frequency trajectories shown in gray (Supplementary Tables 5, 9, 10, 11 and 14).

**Fig. 3.**
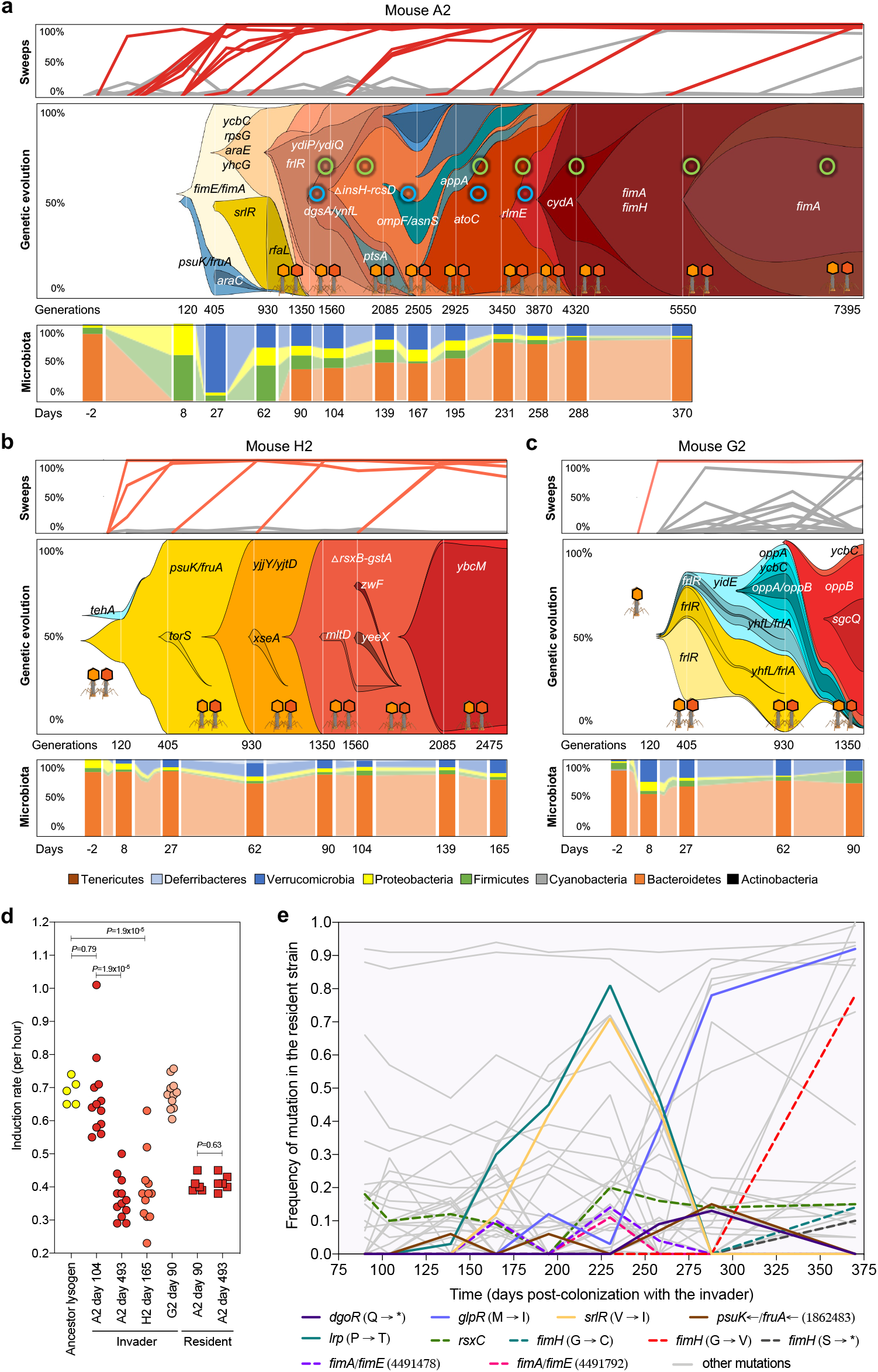
Evolution of invader *E. coli* when co-existing with the resident strain. **a-c,** Frequency of the mutations identified in the invader *E. coli* population across time in mouse A2, H2 and G2 (top panels), the corresponding Muller Plots (middle panels) and microbiota compositions at the phylum level (bottom panels). The spread of *de novo* mutations is indicated by the gene targets where they emerged, intertwined with HGT events, mediated by two phages (in light and dark orange) and two plasmids (green circle – small plasmid: ~69Kb; blue circle – big plasmid: ~109Kb) acquired from the resident strain which started to be detected on day 90 in mouse A2 but much earlier in the other mice. In mouse H2 invader evolution could be followed along 2475 generations, after which extinction occurred and in mouse G2 evolution could be followed for 1350 generations. Mutations that reached frequency above 95% are highlighted in color, other frequency trajectories are shown in gray (Supplementary Tables 5, 8, 12 and 13). **d,** Evidence for prophage domestication, shown by a reduction of lytic induction during gut adaptation. The ancestral lysogen and clones randomly sampled from mouse A2 at days 104 and 493, and mouse H2 at day 165, which bear Nef and KingRac prophages (lysogens), had their prophage induction rate (per hour) measured in a mitomycin C assay. *P* values were calculated by a t-test. **e,** Molecular evolution of the resident strain in mouse A2. The loci highlighted in color were targets of evolution in both the resident and the invader *E. coli* strains. Mutations hitting genes with metabolic functions are indicated with solid lines (*i.e. dgoR, glpR, srlR,psuK/fruA* and *lrp*).

Mouse D2 developed an ecotype characterized by long-term coexistence of invader sublineages within a species rich gut microbiome. No single mutation fixed over ~6540 generations, even though the total population size of the invader *E. coli* was large (~10^7^-10^8^ CFUs/g). The lack of fixations indicates that diversifying selection dominates over directional selection and maintains genetic variation over extended periods. In mice E2 and B2, we also observe signals of diversifying selection and prolonged polymorphism lifetimes. However, the coexistence of ecotypes is only transient and overridden by occasional fixations (in mouse B2 after 6000 generations, in mouse I2 after 1500 generations), in accordance with model results (Good et al., 2018).

The first mutation to arise in the D2 invader *E. coli* was a single-nucleotide polymorphism (SNP) (Supplementary Table 10), lying in the promotor region of *psuK*, which encodes a pseudouridine kinase. This SNP alters *E. coli* growth *in vitro* (Extended Data Fig. 4, Supplementary Table 15). It reached the highest frequency by day 62 (85%), yet failed to reach fixation over more than a full year of evolution in the D2 mouse. Mutations in *dgoR* may have been a driver of diversifying selection in mouse D2 and reduced the sweep rates in other mice (Fig. 2, Supplementary Tables 9, 11 and 14). Specifically, we found signals of negative-frequency-dependent-selection acting on the *dgoR* mutations in mouse D2 and E2 (Extended Data Fig. 5a, b, Supplementary Table 7), consistent with previous results (Ramiro et al., 2020) where *dgoR* KO is advantageous when rare but deleterious at high frequency. Furthermore, competitive fitness assays in mice showed that the fitness effect of a *dgoR* KO depends on the presence of the resident. In one-to-one competition experiments in the mouse gut, the *dgoR* KO mutant decreases to very low frequency when the resident is present but only to ~10% when it is absent (linear mixed model followed by anova test: *dgoR* KO mutant frequency in presence versus absence of resident, P<0.0001; Extended Data Fig. 6, Supplementary Table 16). Evidence of strong selection to consume galactonate is further provided by the observation of recurrent mutations in *dgoR* (Supplementary Table 10), both in our mice and in other work on *E. coli* strains adapting to the gut of streptomycin treated mice, where a resident is absent (Lescat et al., 2017; Ramiro et al., 2020). The combined evidence that disruption of *dgoR* is under negative-frequency-dependent-selection provides a plausible explanation for the long-term maintenance of *E. coli* polymorphism in mouse D2 even in the face of other mutations (Good et al., 2018). This form of selection was also observed in mouse B2 for *srlR*. Here, mutations altering competition for sorbitol (Supplementary Table 9) also have an advantage when rare (~0.4% per generation) but are a disadvantage when common (Extended Data Fig. 5c), consistent with previous findings in streptomycin-treated mice (Lourenço et al., 2016).

Mouse A2, H2, and G2 exhibited evolution under directional selection (Fig. 3). A priori, the presence of a closely related strain in these mice could alter the mode of invader evolution in different ways: by reducing population size and slowing down adaptation, by reducing the number of available nutritional niches (Jousset et al., 2016), or by increasing selection strength and enhancing the rate of fixations. In our experiments, the sweep rate was not correlated with population size, but was significantly higher when co-existence between invader and resident occurred (Welch two-sample t-test t = 2.88, df=4.23, *P-*value = 0.021, one sided). This indicates that the sweep pattern is driven by selection induced by the resident. The resident strain carries prophages that can attack the invader (Frazão et al., 2019), which creates new selection pressures that can alter the course of evolution for the invader *E. coli*. Indeed, co-existence of the resident and invader *E. coli* strains resulted in the co-occurrence of selective sweeps and two distinct mechanisms of HGT by transduction and conjugation (Fig. 3a, Extended Data Fig. 7, Supplementary Tables 5 and 8). During the first thousand generations, when the invader lineage was dominant in mouse A2, seven mutations accumulated and reached high frequency. After this period, and with the concurrent rise in frequency of the resident *E. coli* (Fig. 1a), a strong interlude of HGT marked the evolution of invader *E. coli* in the gut. This occurred initially by phage-mediated transfers, with the rapid formation of a double lysogen, followed by the acquisition of two plasmids from the resident strain (a large plasmid of ~109Kb and a smaller plasmid of ~69Kb) (Fig. 3a). The process of phage-driven HGT was observed in all the mice where invader and resident *E. coli* co-existed (Supplementary Tables 8, 12 and 13, Fig. 3). Interestingly, of the seven putatively active prophages in the resident, only two (Nef and KingRac) transferred to the invader *E. coli* genome during long term co-existence, and the double lysogens that formed rapidly reached high frequencies and were maintained in the gut for a long time.

Conjugation was only detected in mouse A2 and only the small plasmid was maintained for over 7000 generations, being detected in 96% of the clones at day 493 (Extended Data Fig. 7 and Supplementary Table 17). This indicated that any potential benefits that the large plasmid could bring (Valle et al., 2020) were out-weighed by its cost in the invader *E. coli* background. The context dependency of plasmid fitness has also recently been seen across gut isolates (Valle et al., 2020). Overall, 22 selective sweeps accounted for the evolution of invader *E. coli* in mouse A2 for >7000 generations; in mouse H2, seven sweeps were observed in <3000 generations (Fig. 3 and Supplementary Tables 5, 8 and 13).

Phage-driven HGT was a hallmark of the mode of evolution with the resident *E. coli* (Fig. 3). These phages, when fixed, bring metabolic benefits to the invader (Frazão et al., 2019), but induction into a lytic cycle can also entail fitness costs as it leads to the lysogens’ death. *In vitro*, the induction rate for lysogens of the invader *E. coli* was higher than that of the resident strain (Fig. 3d) and *in vivo*, induction rates are likely to be even higher, as it is easier to observe phage-driven HGT in the mice than in the Petri-dish (De Paepe et al., 2016). Theoretical modeling of lysogens with different induction rates predicts that selection should lead to a reduction of the induction costs experienced by the invader *E. coli* during its long term evolution with the resident *E. coli* strain (if the resident has a lower induction rate) (Cortes et al., 2019; Frazão et al., 2019). Clones isolated just after acquisition of the two prophages (at day 104, mouse A2) (Fig. 3a) have a mean induction rate of 0.67 h^-1^ (0.07 2SE), similar to that of a newly formed double lysogen (0.69 h^-1^ (0.02 2SE) (Fig. 3d, Supplementary Table 18). Clones sampled after long-term evolution (at day 493 post-colonization in mouse A2, or day 165 in mouse H2), showed a significantly lower rate of lysogen induction: 0.37 h^-1^ (0.04 2SE) for mouse A2 and 0.39 (0.06 2SE) for mouse H2, (Unpaired t test: *P*=2×10^-7^ and *P*=9×10^-5^, respectively) (Fig. 3d, Supplementary Table 18). Furthermore, the induction rate of the resident *E. coli* clones was maintained through time (0.41 h^-1^ (0.01 2SE)) (Fig. 3d, Supplementary Table 18). These results indicate that under evolution with the resident *E. coli*, lysogens in the invader population adapt over a few thousand generations to avoid the high costs of induction occurring after they are formed and evolve an optimal level of induction. Given that no mutations were detected in the prophage sequences, the observed reduction of lytic induction was likely caused by sweeps that accumulated in the bacterial chromosome. This indicates epistasis between prophages and mutations, as some of the accumulated mutations may be beneficial only in the presence of the prophages. Thus, *E. coli* can ‘domesticate’ its phages to maintain the benefits they bring and avoid the costs of induction into a lytic cycle. This may help explain why it is difficult to induce phages from gut isolates of *E. coli* strains we used in this experiment.

### Evolutionary convergence between *E. coli* strains

Many of the adaptive mutations observed (e.g., frameshifts and missense mutations in *frlR, srlR, dgoR, kdgR*) likely caused changes in the ability of the invader to consume different resources, supporting the notion that adaptation to different resources is important for structuring the microbial diversity in the intestine (Conway and Cohen, 2015; Leatham-Jensen et al., 2012). Given that resource competition can also shape the evolution of the resident *E. coli* (Roughgarden, 1976), we pool-sequenced its clones from mouse A2. Aligning the reads against the reference genome revealed 47 mutations in the resident *E. coli* (Fig. 3e, Supplementary Table 19). Of the mutations observed, 28% occurred in genetic targets that were also seen in the invader *E. coli* strain in mouse A2 and others (Fig. 3e, Supplementary Tables 8 to 14, 19 and 20). Possibly, the *E. coli* strains share similar niches and influence each other’s ability to form stable ecotypes. An example of this effect is detectable in the time series obtained for the two *E. coli* strains in mouse A2: in the invader lineage, the frequency of the *srlR* mutation rose between days 27 and 62, then decreased to undetectable levels by day 90 (Fig. 3a, Supplementary Table 8). Likewise, a mutation in *srlR* was detected in the resident *E. coli* by day 165, reached 71% frequency by day 230, but by day 288 was no longer detected (Fig. 3e, Supplementary Table 19). Similarly, the intergenic region *psuK/fruA* was a target of selection across invader *E. coli* populations isolated from different mice (71%, Supplementary Table 7) and a target in the genome resident *E. coli* of mouse A2 (Fig. 3e). The *glpR* gene was also mutated in the resident of mouse A2 (Fig. 3e, Supplementary Tables 19 and 20) and was a target of selection in invader *E. coli* in another animal (Supplementary Table 9).

Beyond metabolic evolution, mutations in targets related to cell structure were also convergent between invader and resident bacteria. The *fim* operon was recurrently mutated throughout the colonization in both invader and resident strains (Fig. 2b, and Fig. 3a, e). Mutations occurred first in the *fimE/fimA* intergenic region, then in the structural genes *fimA* and/or *fimH* of the invader or the resident. The *fimH* gene is highly polymorphic in *E. coli* (Tenaillon et al., 2010) and an important contributor to *E. coli* pathogenesis (Simms and Mobley, 2008). Changes in the *fim* locus can lead to changes in motility or biofilm formation (Simms and Mobley, 2008). *In vitro* assays showed that an evolved clone, which we sequenced (Supplementary Table 21), presented reduced motility (Extended Data Fig. 8a), but not biofilm capacity (Extended Data Fig. 8b, Supplementary Table 22), and when growing under static conditions formed aggregates of cells at the liquid-air interface (Extended Data Fig. 9). This evolved phenotype is similar to that observed during *in vitro* adaptation of *E. coli* to a fluctuating environment (Behringer et al., 2018) and indicates that the A2 invader lineage may have evolved to occupy a different niche than its ancestor. The changes at the *fim* locus and other loci important for *E. coli* metabolism (Supplementary Tables 8 and 19) indicate evolutionary parallelism occurred in both invader and resident strains. These data illustrate the potential for a high level of evolutionary convergence between two phylogenetically distinct strains colonizing the intestine of the same host.

### Frequency-time statistics of mutational trajectories

To quantitatively characterize the modes of evolution and underlying selection, we used the time-resolved mutation trajectories resolved in Figs. 2 and 3 to estimate the probability that a mutation reaches frequency *x* and the average time it takes to reach frequency *x*. These functions, *G*(*x*) and *T*(*x*), are discussed in Methods and plotted in Fig. 4a, b and Extended Data Fig. 10. Next, we evaluated two simple summary statistics: the probability that a mutation established at an intermediate frequency reaches near-fixation, *p* = G(0.95)/G(0.3), and the ratio of average times to near-fixation and to an intermediate frequency, *τ* = T(0.95)/T(0.3). The threshold frequencies used here are chosen for practical purposes; their details do not matter for the subsequent results. The joint statistics of fixation probabilities and times allow three scenarios of adaptive evolution to be distinguished: diversifying selection leading to (i.) coexistence of ecotypes and directional selection leading to (ii.) clonal interference or (iii.) periodic sweeps (Fig. 4c and Methods). Under predominantly diversifying selection, few established mutations fix and relative fixation times are long (*p* ≪ 1, *τ* ≫ 1). This mode is consistent with the data from mice D2, B2, and E2. Under clonal interference driven by directional selection, only some fraction of the established mutations fixes and relative fixation times are short (*p* < 1, *τ* < 2). This mode was observed in mice A2, G2, and I2. For periodic sweeps to occur, all established mutations fix and the time to near-fixation is about twice the time to half-fixation, given that individual mutations under positive selection follow a sigmoid frequency trajectory (*p* ≈ 1, *τ* ≈ 2). This mode was found in mouse H2 (Fig. 4c). Together, the *p-τ* test infers a dominant mode of evolution occurred in each mouse for which we have long-term data (D2, B2, and A2); the inference is somewhat noisier for the shorter trajectories. These findings are in line with results on *Pseudomonas fluorescens*, where adaptive diversification depends on the level of community diversity (Jousset et al., 2016) and negative-frequency-dependent-selection operates between evolved morphotypes in the absence, but not in the presence of a natural community (Gómez and Buckling, 2013).

**Fig. 4.**
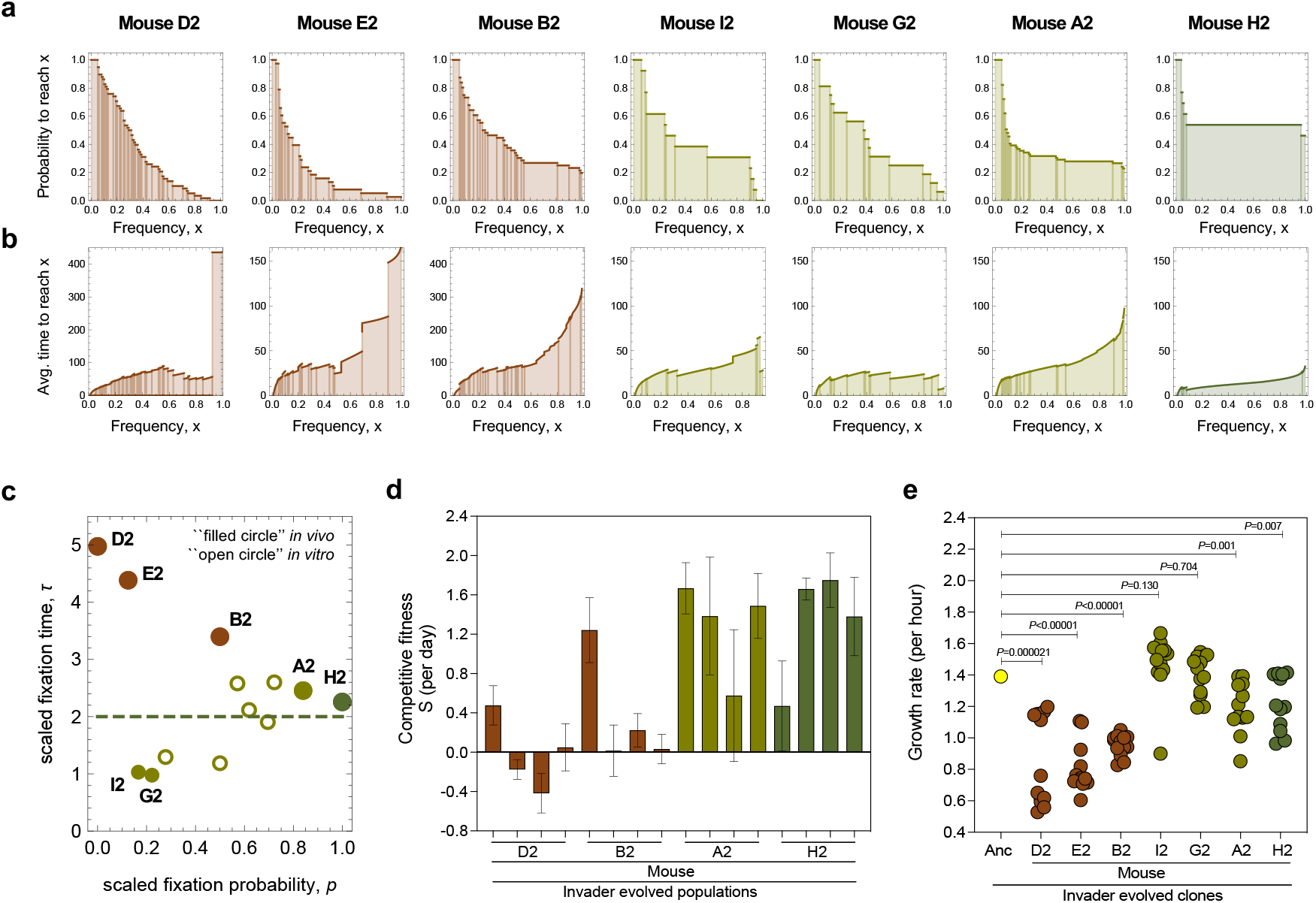
Modes of evolution and fitness tradeoffs. **a-c,** A *p-τ* selection test captures the joint statistics of fixation probabilities a, and times b. This test c, identifies dominant diversifying selection in mice D2, B2, and E2 (brown), and directional selection generating clonal interference in mice A2, G2, and I2 (green) or periodic sweeps in mouse H2 (dark green). In mouse D2, where no fixations occur, we use the time to the end of the experiment lower bound for τ. The reference value *τ* ≈ 2 expected for directional selection in the low-mutation regime is shown as dashed line. Short-term trajectories (< 2000 generations) are marked by small symbols. The data from Lenski’s *in vitro* evolution experiment (Good et al., 2017), for the same lines as in Fig. 1, are shown as open circles. **d,** Host specificity of *in-vivo* adaptation. Relative fitness of pools of evolved clones from mice D2, B2, A2 and H2 when competing against the ancestor in new mice (*n*=4, 2 male and 2 female per competitive fitness assay of each of the independently evolved populations). Error bars represent SE. **e,** *In vivo* adaptation leads to growth trade-offs *in vitro*. Growth rate (per hour) measured in LB medium (3-4 replicates) of randomly isolated invader *E. coli* clones evolved in the gut of mice D2, B2, E2, I2, A2, H2 and G2 during 436, 493, 167, 104, 493, 165 and 90 days, respectively. *P* values were calculated by one sample t-test.

Specific fitness tradeoffs further delineate these *in-vivo* modes of evolution. First, we performed *in vivo* competition experiments between evolved populations and the ancestral clone (Fig. 4d, Supplementary Table 23). Only under directional selection, the evolved populations acquired a net benefit over the ancestral ones. This indicates that when evolution is dominated by diversifying selection the competitive fitness of the populations is more host-specific than when evolution is dominated by directional selection. Second, the populations evolved under diversifying selection show an average 51% reduction of *in vitro* growth compared to the ancestral K12 strain (Fig. 4e, Supplementary Table 24). K12 is a laboratory-adapted strain first isolated from the mammalian gut (Bachmann, 1972). The cost observed here reflects fitness tradeoffs between the laboratory environment and rewilding in the gut. Again, this effect is stronger under predominantly diversifying selection than under directional selection.

The variability of selection regimes of *E. coli in vivo* is higher than in the long-term *in vitro* experiment conducted over the years by Lenski’s team (Good et al., 2017). Here we find a rapid onset of diversifying selection or predominantly directional selection. Of course, more *in vivo* experiments are needed to map conditions determining the relative likelihood of these evolutionary modes. In contrast, all *in-vitro* non-mutator populations show a common pattern of clonal interference under strong directional selection (Fig. 4c and Extended Data Fig. 11); at a comparable speed of molecular evolution (Fig. 1c), diversifying selection emerges only at later times (Good et al., 2017).

## Discussion

In summary, our results offer a longitudinal view of *E. coli* evolution in complex ecosystems over thousands of generations. Metabolic adaptation continuously generates diversifying selection, which can induce the formation of transient ecotypes and the maintenance of functional and genetic diversity within an invader lineage. The dynamics of diversifying selection can be overridden by unconditionally beneficial mutations; an example seen in our experiments is phage-induced directional selection. In this case, we observe ecotype competition between invader and resident *E. coli* strains. How the eco-evolutionary dynamics in the gut plays out on longer time scales and how the relative frequency of the two modes of evolution identified here shape microbiome diversity are important questions for future work.

We conclude that metabolic interactions, by generating ecotypes within and between genetic lineages, shape the mode of evolution of *E. coli* strains inhabiting the mammalian gut. These evolutionary processes, in turn, determine stability, turnover, and diversity of ecotypes.

**Supplementary Information** includes Supplementary Tables.

## Acknowledgments

We thank Jonathan Howard for discussions, Arjan deVisser for a critical reading of the manuscript, and Benjamin Good for sharing data from *in vitro* experiments. We would also like to thank the personnel of the IGC’s Rodent Facility, Genomic Facility and the Bioinformatics Unit for their assistance as well as Roberto Balbontin for the marked resident clone. N.F. was supported by “Fundação para a Ciência e Tecnologia” (FCT), fellowship SFRH/BPD/11075/2015, AK by a cooperation agreement between IGC and the University of Cologne. This work was also supported by project Global Gut Health Nature Research/Yakult Grant 623877 and ONEIDA project (LISBOA-01-0145-FEDER-016417) co-funded by FEEI – “Fundos Europeus Estruturais e de Investimento” from “Programa Operacional Regional Lisboa 2020”, by national funds from FCT, and by Deutsche Forschungsgemeinschaft grant CRC 1310 (to M.L.).

## Author contributions

I.G. and N.F. designed and coordinated the study. N.F. performed the experiments with help from D.G. A.K. performed the bioinformatic analysis. M.L. developed the selection tests. I.G., N.F. and M.L. analysed the results and wrote the manuscript. All authors gave final approval for publication.

## Author Information

The authors have declared no competing interests. Correspondence and requests for materials should be addressed to I.G. or N.F.

## Methods

### *Escherichia coli* clones

The ancestral invader *E. coli* strain expresses a Yellow Fluorescent Protein (YFP), and carries streptomycin and ampicillin resistance markers for easiness of isolation from the mouse feces [galK::amp (pZ12)::PLlacO-1-YFP, strR (rpsl150), ΔlacIZYA::scar]. An *E. coli* strain used for the *in vivo* competition experiments is isogenic to the ancestral invader but expresses a Cyan Fluorescent Protein (CFP) and carries streptomycin and chloramphenicol resistance markers [galK::amp (pZ12)::PLlacO-1-CFP, strR (rpsl150), ΔlacIZYA::scar]. The resident *E. coli* lineage was isolated from the feces along time using McConkey + 0.4% lactose medium, as previously described (Frazão et al., 2019). All the resident clones sampled from each mouse belong to *E.coli* phylogenetic group B (Frazão et al., 2019).The invader *E. coli* strains (YFP and CFP) derive from the K-12 MG1655 strain (DM08) and exhibit a *gat* negative phenotype, *gatZ*::IS1 (Barroso-Batista et al., 2014). The resident *E. coli* clone used for the competition experiments in the mouse gut expresses a mCherry fluorescent protein and a chloramphenicol resistance marker, allowing to distinguish the invader and resident strains in the mice feces.

*E. coli* clones were grown at 37°C under aeration in liquid media Luria broth (LB) from SIGMA — or McConkey and LB agar plates. Media were supplemented with antibiotics streptomycin (100 μg/mL), ampicillin (100 μg/mL) or chloramphenicol (30 μg/mL) when specified.

Serial plating of 1X PBS dilutions of feces in LB agar plates supplemented with the appropriate antibiotics were incubated overnight and YFP, CFP or mCherry-labeled bacterial numbers were assessed by counting the fluorescent colonies using a fluorescent stereoscope (SteREO Lumar, Carl Zeiss). The detection limit for bacterial plating was ~300 CFU/g of feces (Frazão et al., 2019).

### *In vivo* evolution and competition experiments

For both *in vivo* evolution and competition experiments we used the gut colonization model previously established (Frazão et al., 2019). Briefly, mice (*Mus musculus*) supplied by the Rodent Facility at Instituto Gulbenkian de Ciência (IGC) drank water with streptomycin (5 g/L) only for 24h before a 4h starvation period of food and water. The animals were then inoculated by gavage with 100 μL of an *E. coli* bacterial suspension of ~10^8^ colony forming units (CFUs). Mice A2, B2, D2, E2, G2, H2 and I2 were successfully colonized with the invader *E. coli*, while mice C2 and F2 failed to be colonized. Six-to eight-week-old C57BL/6J non-littermate female mice were kept in individually ventilated cages under specified pathogen free (SPF) barrier conditions at the IGC animal facility. Fecal pellets were collected during more than one year (>400 days) and stored in 15% glycerol at −80°C for later analysis. In the competition experiments between the invader ancestral *E. coli* and evolved populations, we colonized the mice using a 1:1 ratio of each genotype, with bacterial loads being assessed and frozen on a daily-basis after gavage.

To assess the impact of the mouse resident *E. coli* in the competitive fitness of *dgoR* we performed one-to-one competitions between the invader ancestral and *dgoR* KO clones. We first homogenized the mice microbiotas by co-housing the animals during seven days. The animals were then maintained under co-housing and given streptomycin-supplemented (5g/L) water during seven days to break colonization resistance and eradicate their resident *E. coli*. At this point, the co-housed mice were removed from the antibiotic-supplemented water for two days. The following day, one group of mice was gavaged with an mCherry-expressing resident *E. coli* (n=3 mice) while the other group (n=3) was not, with all animals being individually caged from this point on and receiving normal water without antibiotic. The day after gavage, all mice were colonized with a mix (1:1) of the invader ancestral and the *dgoR* KO clones, and the bacterial loads were assessed and frozen on a daily-basis.

This research project was ethically reviewed and approved by the Ethics Committee of the Instituto Gulbenkian de Ciência (license reference: A009.2018), and by the Portuguese National Entity that regulates the use of laboratory animals (DGAV - Direção Geral de Alimentação e Veterinária (license reference: 008958). All experiments conducted on animals followed the Portuguese (Decreto-Lei n° 113/2013) and European (Directive 2010/63/EU) legislations, concerning housing, husbandry and animal welfare.

### Microbiota analysis

Fecal DNA was extracted with a QIAamp DNA Stool MiniKit (Qiagen), according to the manufacturer’s instructions and with an additional step of mechanical disruption (Thompson et al., 2015). 16S rRNA gene amplification and sequencing was carried out at the Gene Expression Unit from Instituto Gulbenkian de Ciência, following the service protocol. For each sample, the V4 region of the 16S rRNA gene was amplified in triplicate, using the primer pair F515/R806, under the following PCR cycling conditions: 94 °C for 3 min, 35 cycles of 94 °C for 60 s, 50 °C for 60 s, and 72 °C for 105 s, with an extension step of 72 °C for 10 min. Samples were then pair-end sequenced on an Illumina MiSeq Benchtop Sequencer, following Illumina recommendations. Sampling for microbiota analysis was performed until the microbiota composition stabilized (~1 year after the antibiotic perturbation).

QIIME2 version 2017.11 (Caporaso et al., 2010) was used to analyze the 16S rRNA sequences by following the authors’ online tutorials (https://docs.qiime2.org/2017.11/tutorials/). Briefly, the demultiplexed sequences were filtered using the “denoise-single” command of DADA2 (Callahan et al., 2016), and forward and reverse sequences were trimmed in the position in which the 25th percentile’s quality score got below 20. Diversity analysis was performed following the QIIME2 tutorial (Mandal et al., 2015). Beta diversity distances were calculated through Unweighted Unifrac (Lozupone and Knight, 2005). For taxonomic analysis, OTU were picked by assigning operational taxonomic units at 97% similarity against the Greengenes database version 13 (Greengenes 13_8 99% OTUs (250bp, V4 region 515F/806R)) (DeSantis et al., 2006).

### Whole-genome sequencing and analysis pipeline

DNA was extracted (Wilson, 2001) from *E. coli* populations (mixture of >1000 clones) or a single clone growing in LB plates supplemented with antibiotic to avoid contamination. DNA concentration and purity were quantified using Qubit and NanoDrop, respectively. The DNA library construction and sequencing were carried out by the IGC genomics facility. Processing of raw reads and variants analysis was based on previous work (Barreto et al., 2020). Briefly, sequencing adapters were removed using fastp (Chen et al., 2018) and raw reads were trimmed bidirectionally by 4bp window sizes across which an average base quality of 20 was required to be retained. Further retention of reads required a minimum length of 100 bps per read containing at least 50% base pairs with phred scores at or above 20. BBsplit (Bushnell, 2014) was used to remove likely contaminating reads as explained previously (Barreto et al., 2020). Separate reference genomes were used for the alignment of invader (K-12 (substrain MG1655; Accession Number: NC_000913.2)) and resident (Accession Number: SAMN15163749) *E. coli* genomes. Alignments were performed via three alignment approaches: BWA-sampe version 0.7.17 (Li and Durbin, 2010), MOSAIK version 2.7 (Lee et al., 2014), and Breseq version 0.35.1 (Barrick et al., 2014; Deatherage et al., 2015). While Breseq provides variant analysis in addition to alignment, other variant calling approaches were used to identify putative variation in the sequenced genomes, and to verify data from Breseq. A naïve pipeline (Barreto et al., 2020) using the mpileup utility within samtools (Li et al., 2009) and a custom script written in python was employed. Only reads with a minimum mapping quality of 20 were considered for analysis, and variant calling was limited to bases with call qualities of at least 30. At these positions, a minimum of 5 quality reads had to support a putative variant on both strands (with strand bias, pos. strand / neg. strand, above 0.2 or below 5) for further consideration. Finally, mutations were retained if detected in more than one of the alignment approaches, and if they reached a minimum frequency of 5% at a minimum of one time point sampled. Further simple and complex small variants were considered from freebayes (Garrison and Marth, 2012) with similar thresholds, while insertion sequence movements and other mobile element activity was inferred via ismapper (Hawkey et al., 2015) and panISa (Treepong et al., 2018), as well as Breseq, as previously described (Barreto et al., 2020). All putative variants were verified manually in IGV (Robinson et al., 2017; Thorvaldsdottir et al., 2013). Raw sequencing reads were deposited in the sequence read archive under bioproject PRJNA666769.

### Prophage induction rate

To calculate the maximum prophage induction rate we grew *E. coli* lysogenic clones, starting with the same initial OD600 values: ~0.1 (Bioscreen C system, Oy Growth Curves Ab Ltd), with agitation at 37°C in LB medium in the presence or absence of mitomycin C along time (5 μg/mL) (Frazão et al., 2019). The OD600 values were normalized by dividing the ones in the presence of mitomycin C by the ones in the absence of mitomycin C (sampling interval: 30 minutes). The LN of this ratios along time originates a lysis curve, where the maximum slope corresponds to the maximal prophage induction rate for each clone analyzed. We tested evolved clones from mouse A2, H2 and G2 against the ancestral clone which only carries the Nef and the KingRac prophages. We also tested clones of the resident strain that had evolved in the presence of the invader for more than 400 days (these clones were sampled from mouse A2).

### *E. coli* growth rate, growth curves, cell aggregation, biofilm and motility capacity

To calculate the maximum bacterial growth rate, we grew *E. coli* lysogenic clones, starting with the same initial OD600 values: ~0.1 (Bioscreen C system, Oy Growth Curves Ab Ltd), with agitation at 37°C in LB medium along time using reading intervals of 30 minutes. The LN of the OD600 values along time originates a growth curve, where the maximum slope corresponds to the maximum bacterial growth rate for each clone analyzed.

To test for metabolic differences of the *psuK/fruA* mutation, growth curves of evolved lysogenic *E. coli* clones, bearing the Nef and KingRac prophages, with or without the *psuK/fruA* mutation were performed with the same initial OD600 value (~0.03) for each clone. The clones were grown in glucose (0.4%) minimal medium (MM9-SIGMA) with or without pseudouridine (80 μM) and absorbance values were obtained using the Bioscreen C apparatus during 12 hours.

Frozen stocks of *E. coli* clones were used to seed tubes with 5 mL of liquid LB. These were incubated overnight at 37 °C under static conditions to assess the formation of cell flocks/clumps, observable to the naked eye, in order to evaluate the formation of cell aggregates. Biofilm was tested according a previously published protocol (Beloin et al., 2008) and to evaluate the motility capacity we adapted the protocol from Croze and colleagues (Croze et al., 2011). Briefly, overnight *E. coli* clonal cultures grown with agitation at 37 °C in 5mL LB medium supplemented with streptomycin (100 ug/mL) were adjusted to the same absorbance and a 3uL volume was dropped on top of soft agar (0.25%). Plates were incubated at 37°C and photos were taken at day 1, 2 and 5 post-inoculation to assess swarming motility phenotype.

### Number of *E. coli* generations during mouse gut colonization

To estimate the number of generations of *E. coli* in the mouse gut, we used a previously described protocol to measure the fluorescent intensity of a probe specific to *E. coli* 23S rRNA (as a measure of ribosomal content) that correlates with the growth rate of the bacterial cells (Barroso-Batista et al., 2015). We measured the number of generations of the ancestral *E. coli* clone while colonizing the gut of 2 mice, treated during 24 hours with streptomycin (5 g/L) before gavage, during 25 days.

### Plasmid DNA extraction and PCR detection of ~69Kb (*repA*) and ~109Kb (*repB*) plasmids

Plasmid DNA was extracted from overnight cultures using a Plasmid Mini Kit (Qiagen), according to the manufacturer’s guidelines. Specific primers for the amplification of *repA* and *repB* genes, were used to determine the frequency of the 68935 bp (~69 Kb) and 108557 bp (~109 Kb) plasmids, respectively, in the invader *E. coli* population.

The primers used for *repA* gene were: repA-Forward: 5’−CAGTCCCCTAAAGAATCGCCCC-3’ and repA-Reverse: 5’-TGACCAGGAGCGGCACAATCGC-3’.

For *repB* the primer sequences were: repB-Forward: 5’-GTGGATAAGTCGTCCGGTGAGC-3’ and repB-Reverse: 5’-GTTCAAACAGGCGGGGATCGGC3’.

PCR amplification of plasmid-specific genes was performed in 12 isolated random clones from mouse A2 at days 104 and 493. PCR reactions were performed in a total volume of 25 μL, containing 1 μL of plasmid DNA, 1X Taq polymerase buffer, 200 μM dNTPs, 0.2 μM of each primer and 1.25 U Taq polymerase. PCR reaction conditions: 95°C for 3 min, followed by 35 cycles of 95°C for 30 s, 65°C for 30 s and 72°C for 30 s, finalizing with 5 min at 72°C. DNA was visualized on a 2% agarose gel stained with GelRed and run at 160 V for 60 min.

### Construction of the *dgoR* KO mutant

P1 transduction was used to construct a *ΔdgoR* mutant (*dgoR* KO). This KO strain was created by replacing the wild-type *dgoR* in the invader ancestral YFP-expressing genetic background by the respective knock-out from the KEIO collection, strain JW5627 (Baba et al., 2006), in which the *dgoR* sequence is replaced by a kanamycin resistance cassette. The presence of the cassette was confirmed by PCR using primers dgoK-F: GCGATGTAGCGAGCTGTC, and yidX-R: GGGAATAAACCGGCAGCC. PCR reactions were performed in a total volume of 25 μL, containing 1 μL of DNA, 1X Taq polymerase buffer, 200 μM dNTPs, 0.2 μM of each primer and 1.25 U Taq polymerase. PCR reaction conditions: 95°C for 3 min, followed by 35 cycles of 95°C for 30 s, 65°C for 30 s and 72°C for 30 s, finalizing with 5 min at 72°C. DNA was visualized in a 2% agarose gel stained with GelRed and run at 160 V for 60 min.

### Statistical analysis

Correlation between microbiota diversity measures and *E. coli* loads was performed in R using the statistical package rmcorr (Bakdash and Marusich, 2017). The rate of accumulation of new ISs *in vivo* was compared using Wilcoxon paired signed ranked test for expected and observed insertions, while the rate of selective sweeps correlation was performed using the Spearman Correlation test. Selective sweeps were taken to be mutations or HGT events that reached >95% frequency in the population and kept high frequency until the end of the colonization. Statistical analysis of prophage induction as well as biofilm levels was performed using the Mann-Whitney test in GraphPad Prism (version 8.4.3). A single sample T-Test was used test if the growth rate of evolved invader clones deviates from the mean of the ancestral. P values of <0.05 were considered significant.

Pearson correlation tests between the frequency and the change in frequency of a mutation were performed to search for evidence of negative frequency dependent selection. These were conducted for every mutation that showed parallelism and for each mouse, provided that the mutation was detected in at least four time points. The correlations were calculated in R with cor.test, used for association between paired samples.

Linear mixed models (R package nlme, v3.1 (J Pinheiro et al., 2019)) were used to analyze the temporal dynamics of the *dgoR* KO mutant frequency in the presence or absence of the resident *E. coli*. The frequency of the *dgoR* KO mutant was log10 transformed to meet the assumptions of parametric statistics.

### Statistics of time-resolved mutation frequency trajectories

The following statistics are designed to infer the prevalent type of selection from time-resolved mutant frequency data. Specifically, we use such data to discriminate adaptive evolution under directional selection, which can take place by periodic sweeps or by clonal interference, and adaptation under diversifying selection (to be defined below).

We use two test statistics for frequency trajectories of established mutants (i.e., mutants that have overcome genetic drift):

- the frequency propagator *G*(*x*), defined as the probability that a trajectory reaches frequency *x*
- the sojourn time *T*(*x*), defined as the time between origination at a threshold frequency *x*_0_ and the first occurrence at frequency *x*, averaged over all trajectories reaching frequency *x*. In terms of the underlying coalescent, *T*(*x*) is the time to the last common ancestor for a genetic clade of frequency *x*.

These observables discriminate the following modes of adaptive evolution:

- **Periodic selective sweeps under uniform directional selection.** This mode is characteristic of simple adaptive processes in small populations, where adaptive mutations are rare enough to fix independently (Desai et al., 2007; Gerrish and Lenski, 1998; Perfeito et al., 2007). Almost all established adaptive mutations reach fixation, and sojourn times to an intermediate frequency *x* > *x*_0_ are of order of their inverse selection coefficient (up to logarithmic corrections):

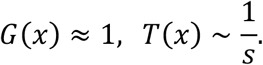
- **Clonal interference under uniform directional selection.** This mode occurs in asexual populations when adaptive mutations become frequent enough to interfere with one another (Desai et al., 2007; Gerrish and Lenski, 1998; Perfeito et al., 2007). Only a fraction of the established adaptive mutations reaches fixation; sojourn times to intermediate frequencies are set by a global coalescence rate 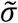 that is higher than the typical selection coefficient of individual mutations (Neher and Hallatschek, 2013):

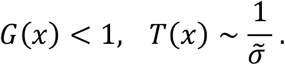 Details of these dynamics depend on the spectrum of selection coefficients and on the overall mutation rate, which set the strength of clonal interference. For moderate interference, where a few concurrent beneficial mutations compete for fixation, we expect a roughly exponential drop of the frequency propagator, *G*(*x*) ~ exp(-*λx*), reflecting the probability that a trajectory reaches frequency *x* without interference by a stronger competing clade. Moderate interference generates an effective neutrality for weaker beneficial mutations and at higher frequencies (Schiffels et al., 2011). This regime has been mapped for influenza (Strelkowa and Lässig, 2012). In the asymptotic regime of a travelling fitness wave, where many beneficial mutations are simultaneously present, the fate of a mutation is settled in the range of small frequencies; that is, at the tip of the wave (Good et al., 2012). In this regime, emergent neutrality affects the vast majority of beneficial mutations and most of the frequency regime (Rice et al., 2015). Hence, the frequency propagator rapidly drops to its asymptotic value *G*(*x* = 1) ≪ 1.
- **Adaptation under diversifying selection.** More complex selection scenarios involve selection within and between ecotypes, i.e., subpopulations occupying distinct ecological niches (Cohan, 2006; Rainey and Travisano, 1998). An important factor generating niches and ecotypes is the differential use of food and other environmental resources. In this mode, ecotype-specific, conditionally beneficial mutations reach intermediate frequencies after a time given by their within-ecotype selection coefficient *s*, but fixation can be slowed down or suppressed by diversifying (negative frequency-dependent) cross-ecotype selection (Good et al., 2018),

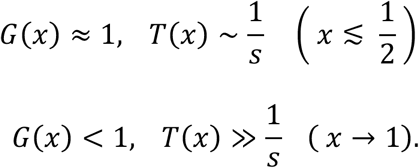 The details depend on the details of the eco-evolutionary model (synergistic vs. antagonistic interactions, carrying capacities, amount of resource competition vs. explicitly frequency-dependent selection). In a model with directional selection within ecotypes, conditionally beneficial mutations rapidly fix within ecotypes, but lead only to finite shifts of the ecotype frequencies. In the simplest case, the resulting dynamics of ecotype frequencies is diffusive, resulting in an effectively neutral turnover of ecotypes (Good et al., 2018). Given negative frequency-dependent selection between ecotypes, fixations become even rarer and can be completely suppressed; that is, ecotypes can become stable on the time scales of observation. The separation of time and selection scales between intra- and cross-ecotype frequency changes is expected to be a robust feature of ecotype-dependent selection: sojourn of adaptive alleles to intermediate frequencies is fast, fixation is slower and rarer. In other words, ecotype-dependent selection is characterized by two regimes of coalescence times *T*(*x*).

Frequency propagators and the coalescence time spectra expected under these evolutionary modes are qualitatively sketched in Extended Data Fig. 10. For periodic sweeps under directional selection (dark green, left column), *G*(*x*) depends weakly on *x* and *T*(*x*) is set by rapid sweeps for all *x*. For clonal interference under directional selection (green, center column), *G*(*x*) decreases substantially with increasing *x* and *T*(*x*) becomes uniformly shorter. Under negative frequency-dependent selection (brown, right column), *G*(*x*) decreases substantially with increasing *x*, while *T*(*x*) substantially increases for large *x* and diverges in case of strong frequency-dependent selection generating stable ecotypes (dashed lines).

### The *p-τ* selection test

This test is based on qualitative characteristics of the functions *G*(*x*), *T*(*x*) and does not depend on details of the evolutionary process. We evaluate *G*(*x*) and *T*(*x*) for host-specific families of frequency trajectories; sojourn times are counted from an initial frequency *x*_0_ = 0.01. Origination times at this frequency are inferred by backward extrapolation of the first observed trajectory segment; the reported results are robust under variations of the threshold *x*_0_ and the extrapolation procedure. We then compute two summary statistics: the probability *p* that a mutation established at an intermediate frequency *x_m_* reaches near-fixation at a frequency *x_f_*,

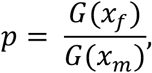

and the corresponding fraction of sojourn times,

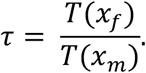

Here we use *x_m_* = 0.3 and *x_f_* = 0.95 to limit the uncertainties of empirical trajectories at low and high frequency; however, the selection test is robust under variation of these frequencies. We find evidence for different modes of evolution:

- The long-term frequency trajectories of mice B2, D2 and E2 are consistent with predominantly frequency-dependent selection (Fig. 2, Fig. 4a-c). The propagator *G*(*x*) is a strongly decreasing function of *x*, resulting in fixation probabilities *p* < 0.5. Sojourn times *T*(*x*) show two regimes with a stronger increase in the frequency range *x >* 0.6, as measured by time ratios *τ* > 3.
- The trajectories of mice A2, G2, and I2 show a signature of recurrent selective sweeps and clonal interference under uniform directional selection (Fig. 4a-c). The propagator *G*(*x*) is a decreasing function of *x*, resulting in fixation probabilities *p* = 0.2 – 0.8, depending on the strength of clonal interference. Fixation times are short, giving time ratios *τ* ≲ 2.
- The shorter trajectory of mouse H2 signals periodic sweeps under uniform directional selection (Fig. 3, Fig. 4a-c). The origination rate of mutations is lower than in the longer trajectories, and *G*(*x*) shows a weak decrease with *p* = 1. Sojourn times *T*(*x*) are short and grow uniformly with *x*, resulting in a time ratio *τ* = 22.5. This pattern is expected under directional selection in the low mutation regime: *T*(*x*) = log[%/(1 – %)]/s for individual mutations with a uniform selection coefficient *s*, leading to *τ* = 2.0 for *x_m_* = 0.3 and *x_f_* = 0.95 (this value is marked as a dashed line in Fig. 4c).
- The trajectories of non-mutator lines in the long-term *in vitro* evolution experiment of Good et al (Good et al., 2017), evaluated over the first 7500 generations, show an overall signal of clonal interference under uniform directional selection (Fig. 4c, Extended Data Fig. 11). The frequency propagators *G*(*x*) are strongly decreasing functions of *x* and sojourn times *T*(*x*) grow uniformly with *x*. We find *p* = 0.2 – 0.8 and *τ* ≲ 2, similar to the pattern in mice A2, G2, and I2.

## Extended Data Figures

**Extended Data Figure 1.**
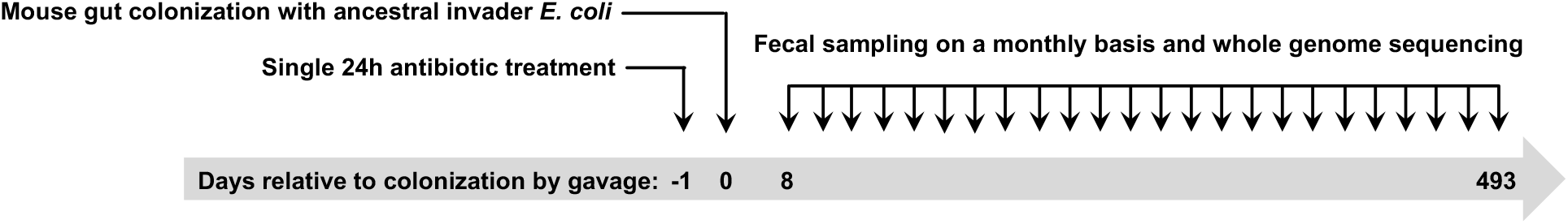
Experimental setup for *in vivo* Long-term Evolution Experiment (*in vivo* LTEE). Independently caged mice (n=7) were treated a single time during 24 hours with streptomycin (5 g/L) in drinking water. Afterwards the antibiotic was removed and colonization of the mice gut was performed by gavage with a suspension of ~10^8^ colony forming units (CFUs) of streptomycin-resistant, yellow-fluorescent protein (YFP) *E. coli* ancestral invader clone. Fecal pellets were collected for 493 days post-gavage (~monthly basis) and stored in 15% glycerol at −80°C for later analysis.

**Extended Data Figure 2.**
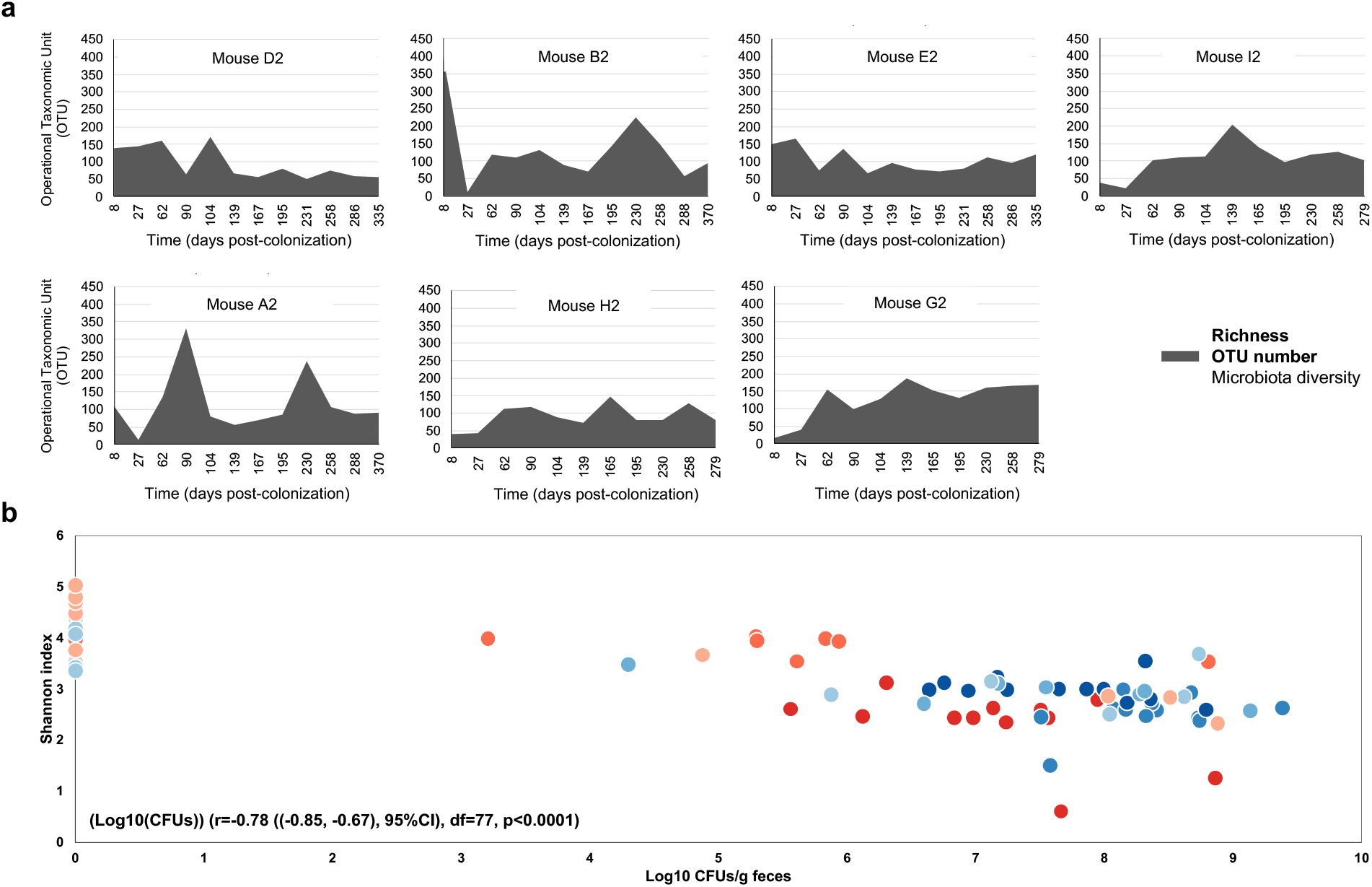
Microbiota diversity metrics. **a,** Richness of the microbiota in mice colonized with the invader E. coli (mice A2, B2, E2, I2, A2, H2 and G2). **b,** Negative correlation between the microbiota Shannon index and the invader *E. coli* persistence among the 9 mice of the experiment (Log10(CFUs)) (r=−0.78 ((−0.85, −0.67), 95%CI), df=77, p<0.0001) (Supplementary Table 2). Each colored circle represents a mouse. P-value was calculated by a repeated measures correlation.

**Extended Data Figure 3.**
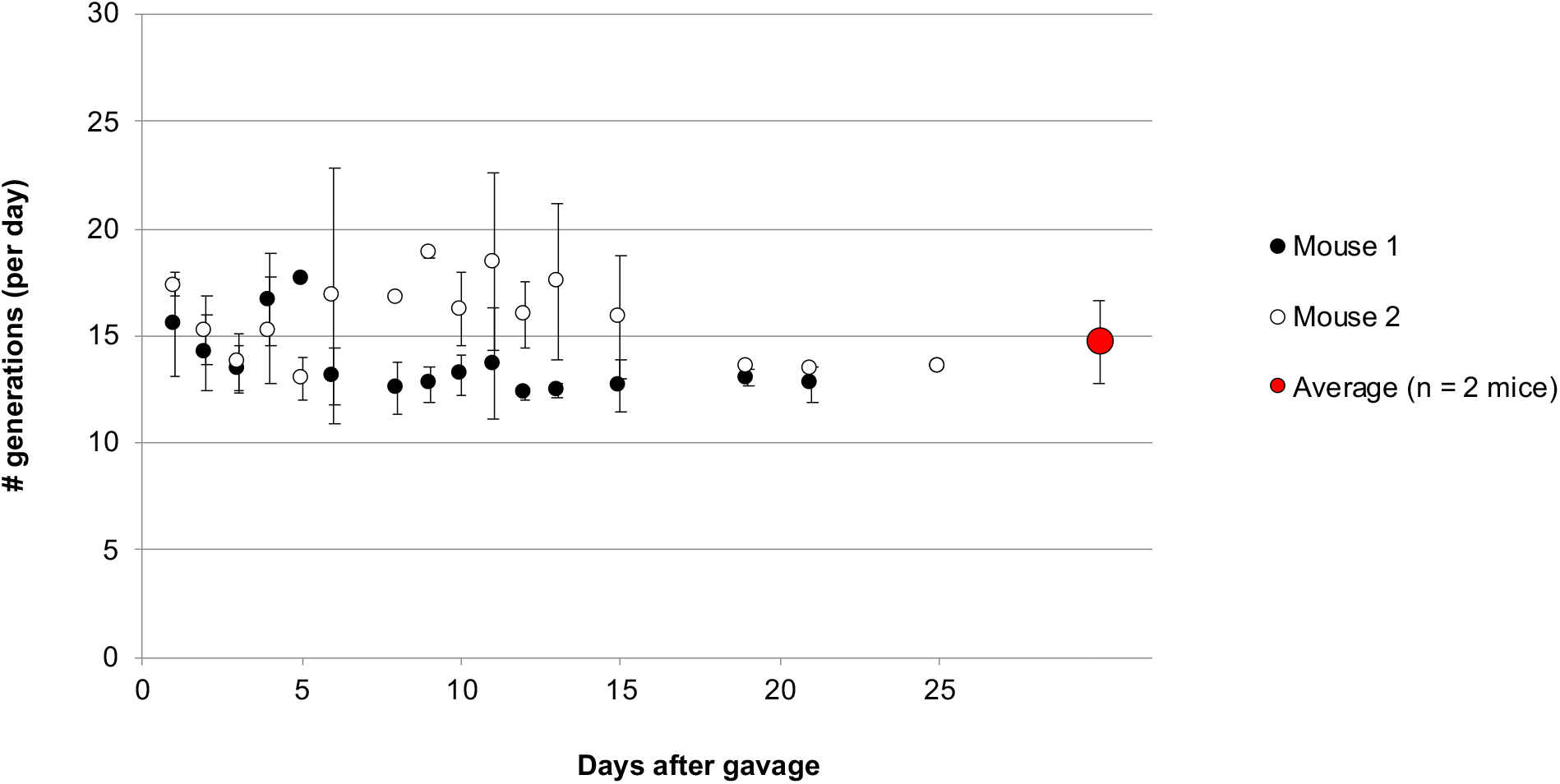
*E. coli* growth rate (generations per day) when colonizing the gut of mice treated with a single 24h streptomycin treatment (5g/L) before gavage. A previously used protocol based on ribosome content allowed us to assess the number of generations of the invader *E. coli* in the mouse gut (see Supplementary Materials). Error bars represent Standard Deviation (SD) (Supplementary Table 4).

**Extended Data Figure 4.**
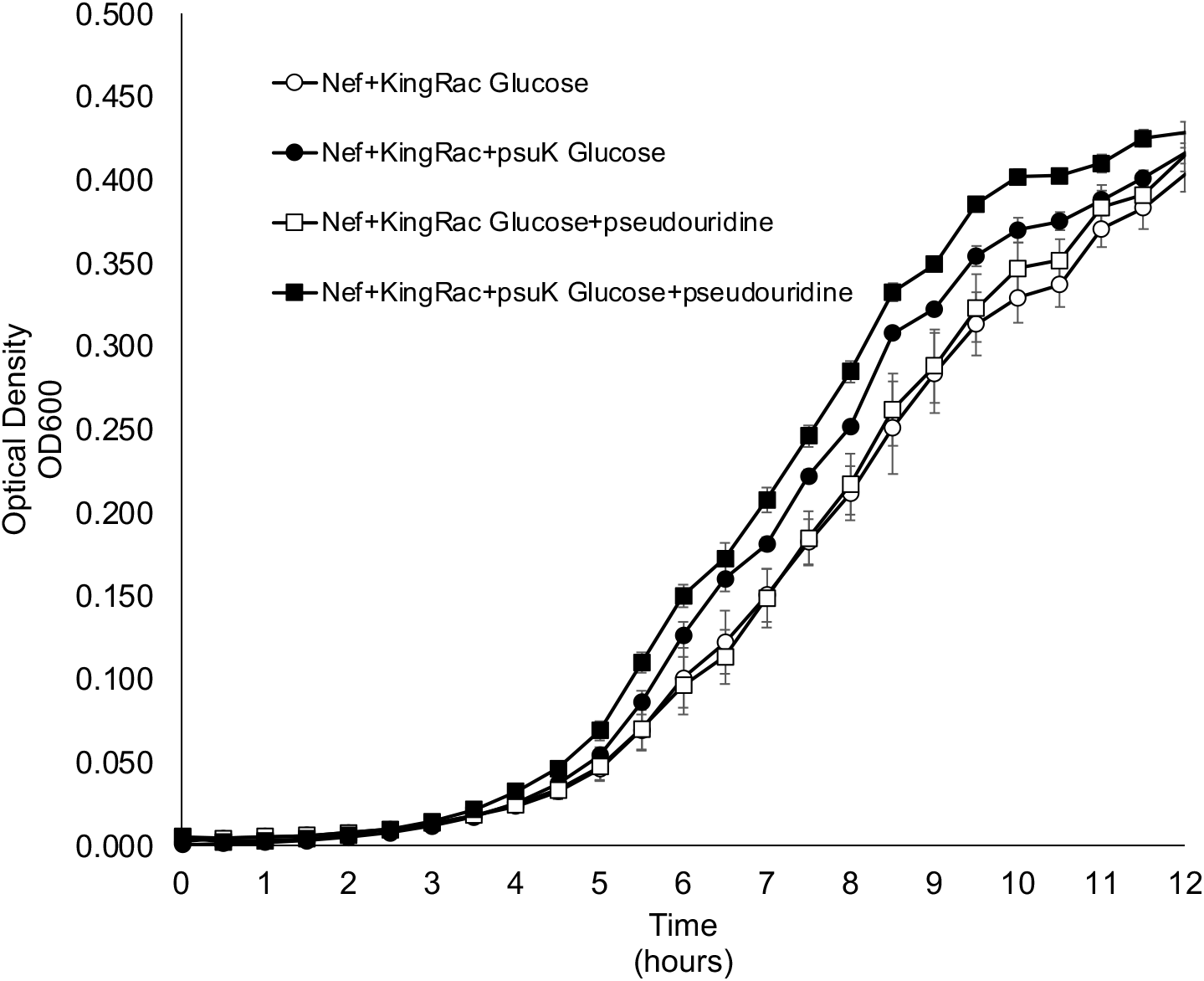
Growth curves of evolved clones. The growth of a double lysogen, carrying the prophages Nef and KingRac (Nef+KingRac), was compared to that of a clone carrying the two prophages and the *psuK/fruA* mutation (Nef+KingRac+*psuK*) *in vitro* (n=6 biological replicates per clone). Nef+KingRac+*psuK* has a higher maximum growth rate than Nef+KingRac (0.80 (0.04 SE) *vs*. 0.69 (0.02 SE), T-test P=0.00036) when grown in glucose (0.4%). When grown in glucose (0.4%) supplemented with pseudouridine (80μM), Nef+KingRac+*psuK* also has a significantly higher maximum growth rate than Nef+KingRac (0.79 (0.02 SE) *vs*. 0.63 (0.04 SE), T-test P=0.0056). Error bars represent Standard Error (SE) (Supplementary Table 15).

**Extended Data Figure 5.**
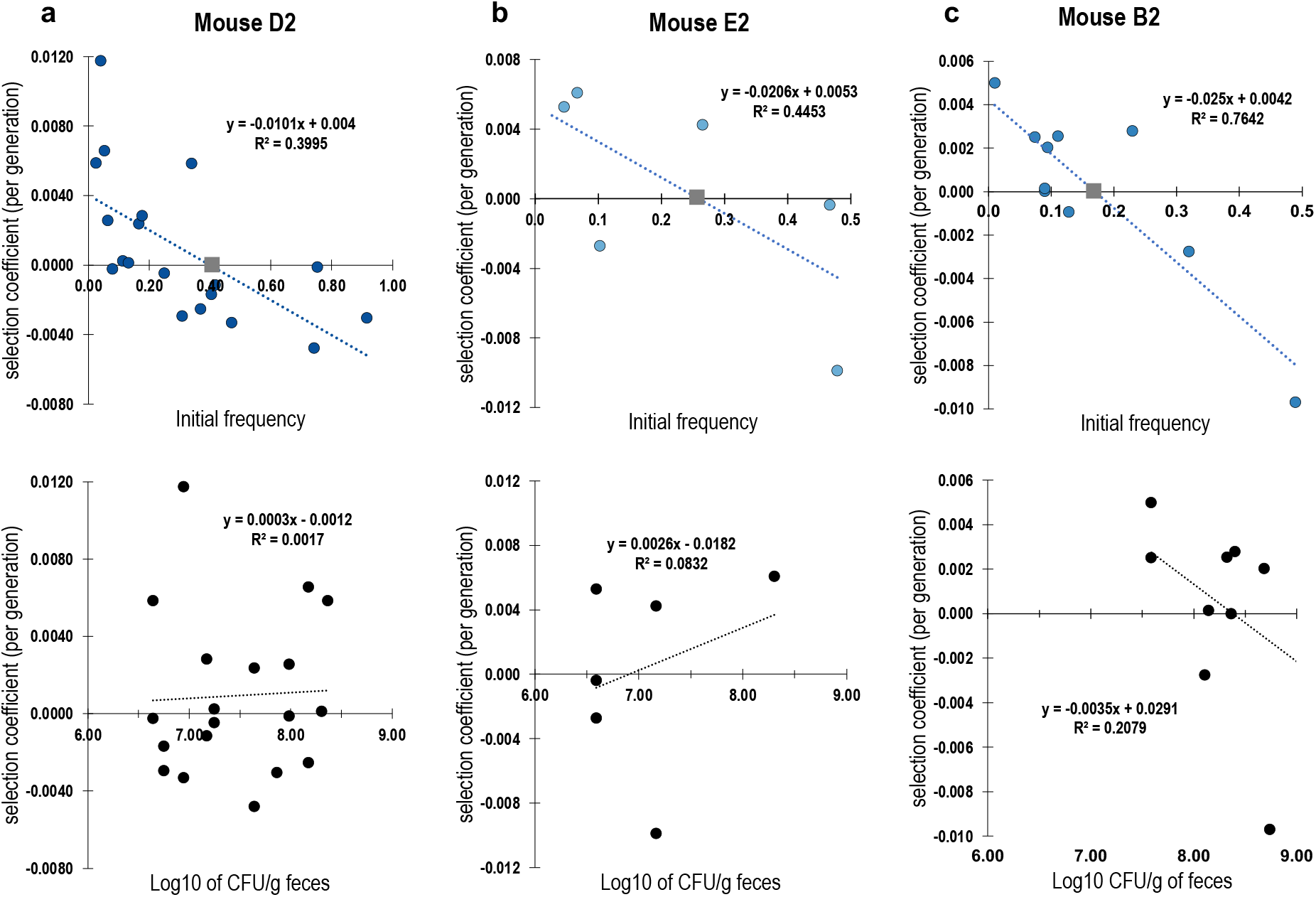
Evidence for negative frequency dependent selection in the frequency trajectory of *dgoR* mutations in mouse D2 and E2 and of *srlR* mutations in mouse B2. The change in the selection coefficient (*s* per generation), measured as the slope between Ln[frequency/(1-frequency)] of each allele at consecutive time points after the mutation was detected in the invader *E. coli* isolated from the feces of a mouse. A change in sign, from a positive *s* at lower frequencies to a negative *s* at higher frequencies, is a signal of negative-frequency dependent selection. Such form of selection should lead to a stable equilibrium frequency indicated by a grey square in the plots, if no other mutations would occur. We note that some forms of density dependent selection could also lead to correlations with initial frequency, however no correlation between *s* and population size (bottom panel), as measured from the CFUs/g of feces, was detected in mouse D2, and a very weak correlation in mouse B2. **a,** In mouse D2, six different alleles were detected in *dgoR*. **b,** In mouse E2, four different alleles were detected in *dgoR*. **c,** In mouse B2, one single allele was detected in *srlR*. All alleles detected were used to calculate the correlations shown in the plots.

**Extended Data Figure 6.**
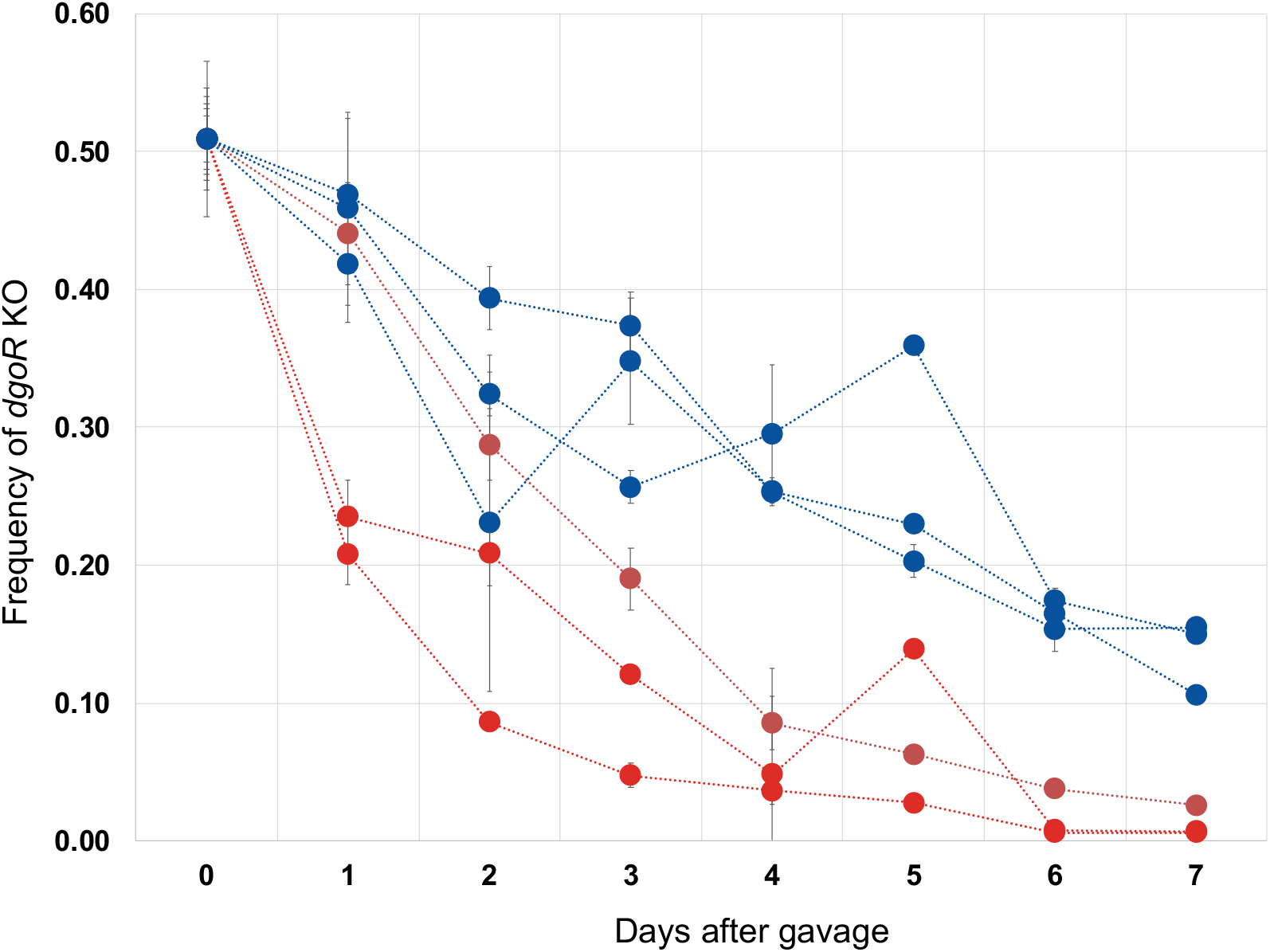
Frequency of the invader *E. coli dgoR* KO mutant when competing with ancestral in the mouse gut in the presence (red colour) or absence (blue colour) of the resident *E. coli* lineage. The presence of the resident (red colour) leads to a significant frequency decrease of the *dgoR* KO mutant, when compared with the resident’s absence (blue colour) (linear mixed model followed by anova test: *dgoR* KO mutant frequency in presence versus absence of resident lineage, *P*<0.0001) (Supplementary Table 16).

**Extended Data Figure 7.**
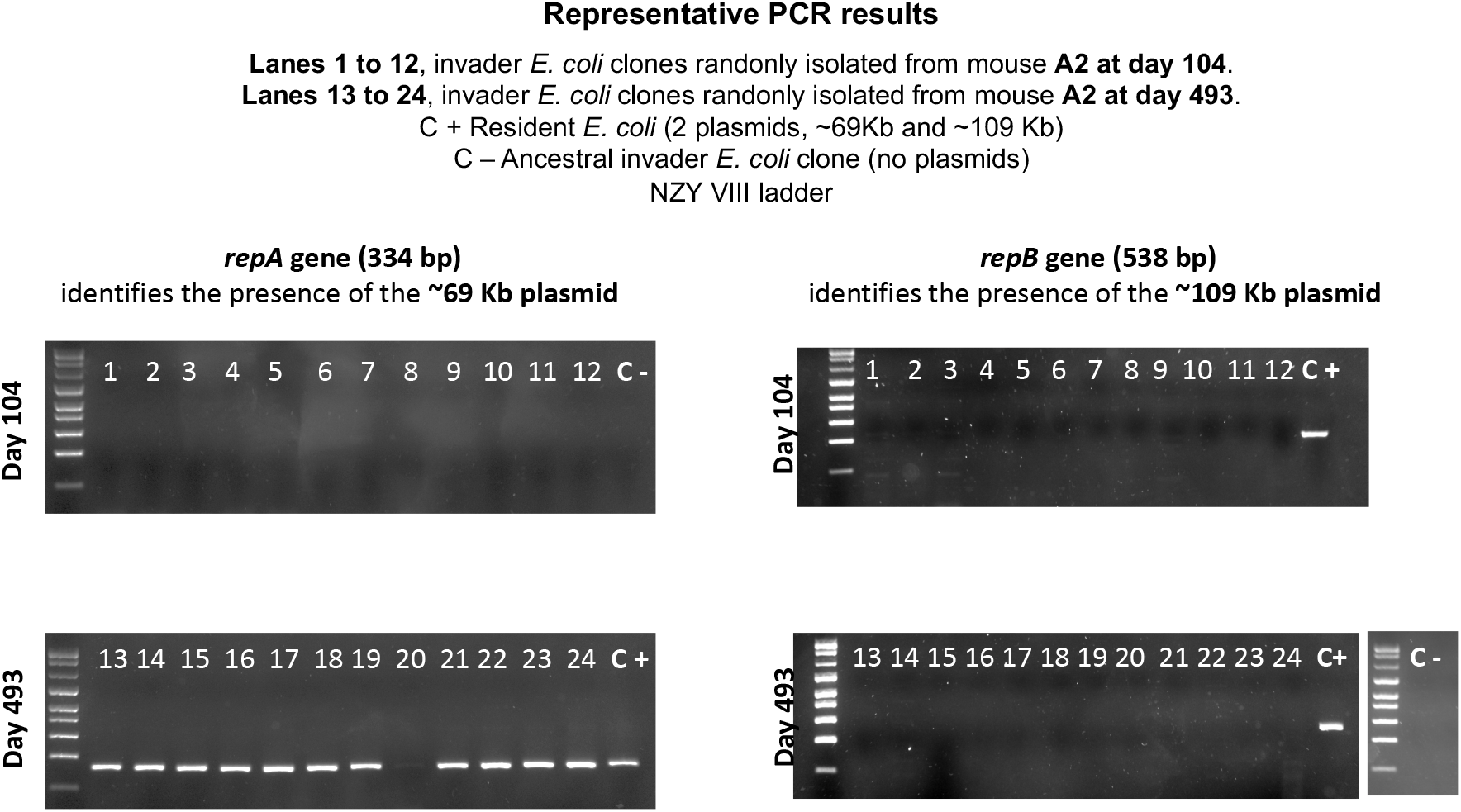
PCR-based plasmid frequency in the invader *E. coli* lineage isolated from the gut of mouse A2. The 334 bp portion of gene *repA*, that detects the ~69 Kb plasmid, inherited from the resident *E. coli* lineage was only detected at day 493 (7395 generations) being at 96% frequency (in 23 out of 24 clones isolated from mouse A2). The *repB* gene (identifying the ~109 Kb plasmid) was never detected in any of the clones isolated either at day 104 or 493.

**Extended Data Figure 8.**
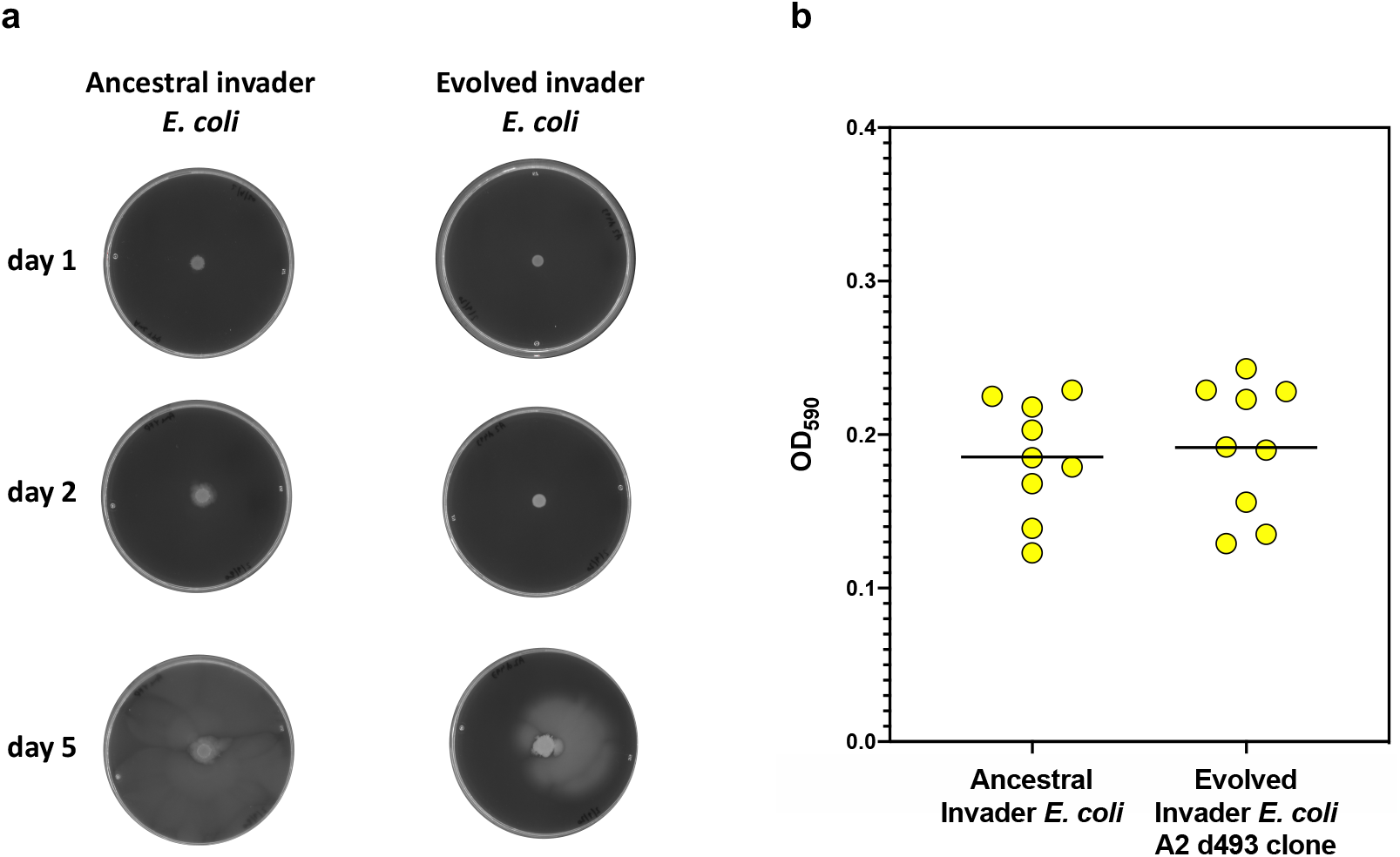
Gut adaptation appears to alter the motility while maintaining the biofilm capacity. **a,** Motility assays (ancestor and evolved clone isolated from mouse A2 at day 493). Each clone drop contained the same number of cells (OD600=0.1 – Bioscreen measurement). Images are representative of the observed phenotype. Three biological replicates were performed per clone. **b,** Biolfilm assay. Three biological replicates per clone, each with 3 technical replicates, were performed (Supplementary Table 22).

**Extended Data Figure 9.**
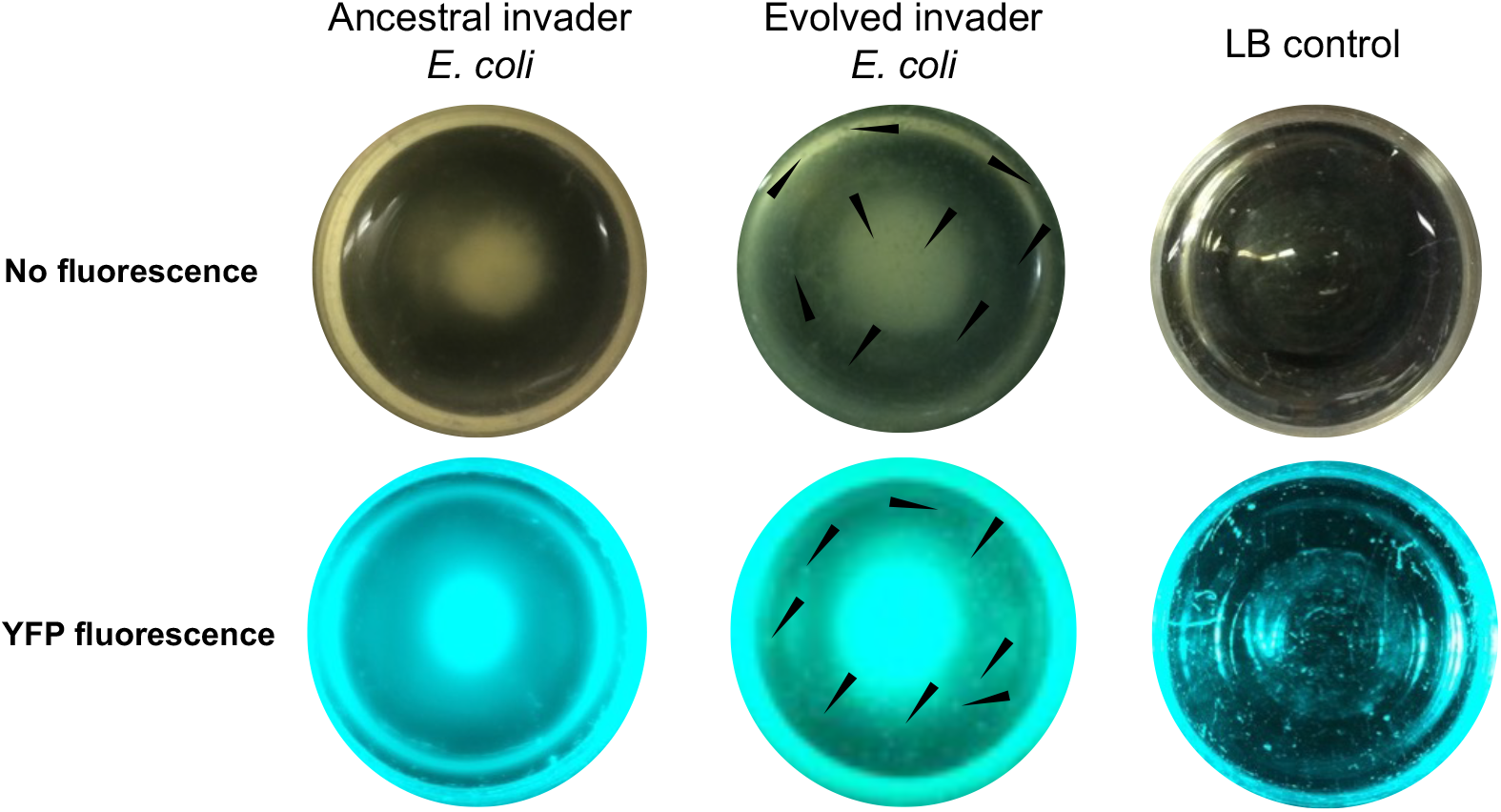
Evolved invader *E. coli* forms aggregates when growing in static conditions. Images are representative of the observed phenotype. Black arrow point to cell aggregates. Three biological replicates per clone, each with three technical replicates, were performed. Ancestral and evolved (isolated from mouse A2 at day 493) clones express a Yellow Fluorescent Protein (YFP) marker. Blue/green LED transilluminator (FastGene) allowed verification that the bacteria were YFP fluorescent and not contaminants.

**Extended Data Figure 10.**
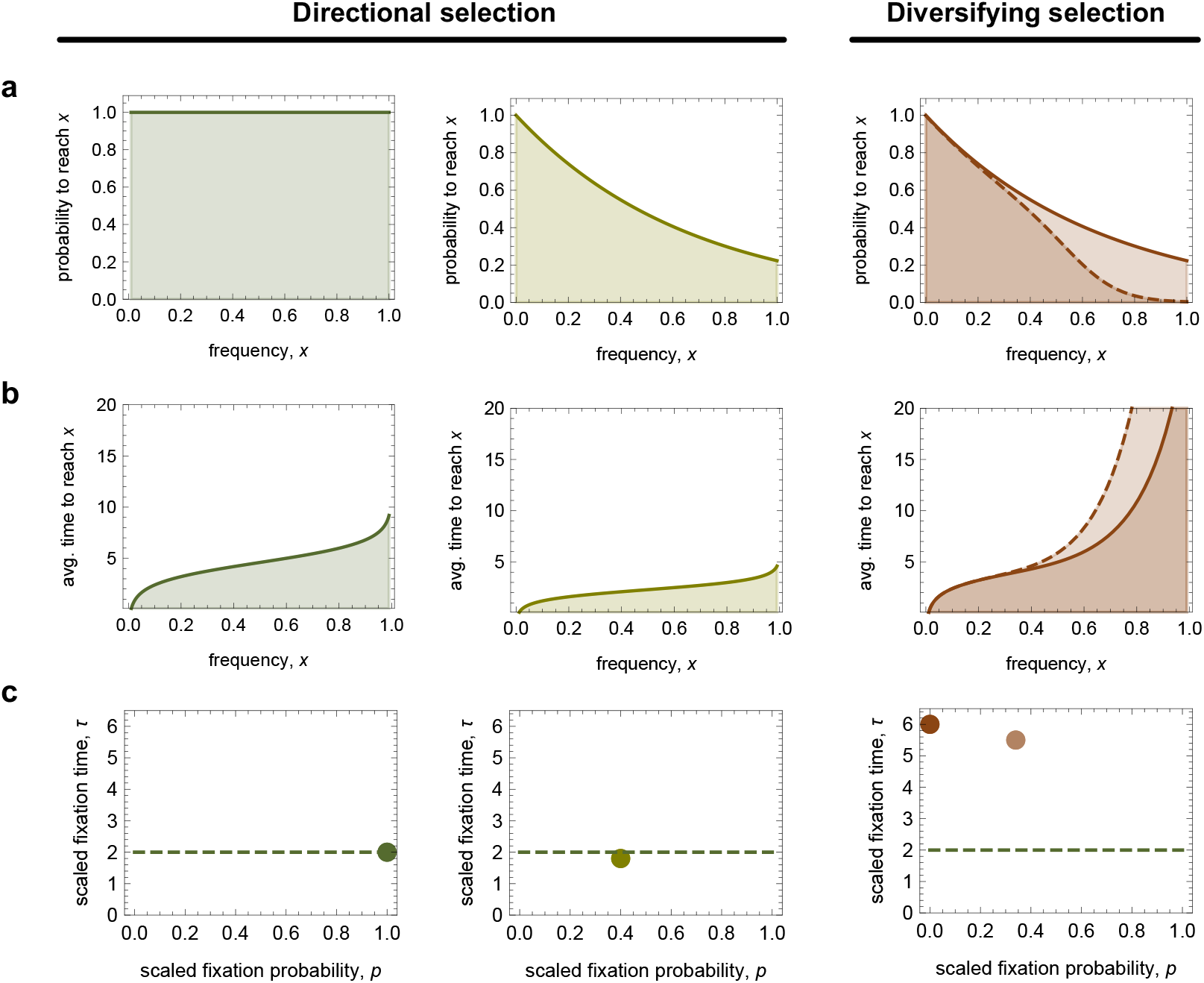
a, Frequency propagator. b, Coalescence time spectrum. c, *p* – *τ* summary statistics under different modes of adaptive evolution. **Periodic sweeps under directional selection (left):** high fixation probability and fast fixation of established, uniformly beneficial mutations. **Clonal interference under directional selection (center):** decreased fixation probability, but even faster fixation of established, uniformly beneficial mutations. **Diversifying selection (right):** decreased fixation probability and slower fixation of established mutations with ecotype-specific fitness advantage. Fixations can be completely suppressed in case of strongly negative frequency-dependent selection across ecotypes (dashed lines); in that case, the total observation time of a mutation trajectory serves as a lower bound for the fixation time in the *p* – *τ* summary statistics. Times are measured in units of the inverse selection coefficient of individual sweeps.

**Extended Data Figure 11.**
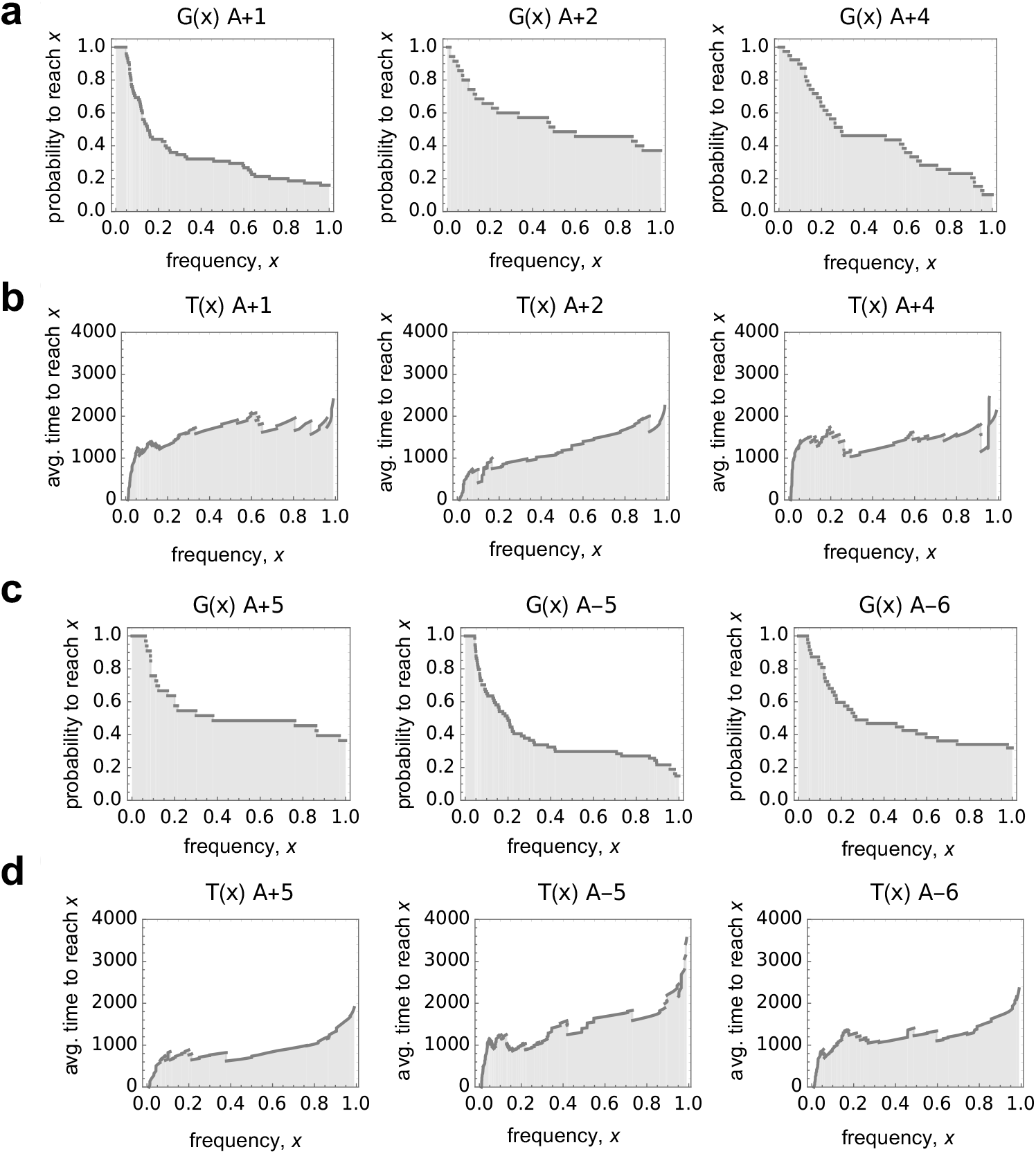
Statistics of *in-vitro* trajectories. Frequency propagators **(a, c)** and coalescence time spectrum **(b, d)** observed in the 6 Lenski populations that did not evolve a mutator phenotype over a period of 7500 generations (populations A+1, A+2, A+4, A+5, A-5 and A-6). Only mutations that reached at least 5% frequency where counted. Time is measured in generations. Under the conditions of the experiment 6.7 generations pass per day.

